# Genomic variation and population histories of spotted (*Strix occidentalis*) and barred (*S. varia*) owls

**DOI:** 10.1101/2020.02.18.954685

**Authors:** Naoko T. Fujito, Zachary R. Hanna, Michal Levy-Sakin, Rauri C. K. Bowie, Pui-Yan Kwok, John P. Dumbacher, Jeffrey D. Wall

## Abstract

Spotted owls (SO, *Strix occidentalis*) are a keystone species inhabiting old-growth forests in Western North America. In recent decades, their populations have declined due to ongoing reductions in suitable habitat caused by logging, wildfires, and competition with the congeneric barred owl (BO, *Strix varia*). The northern spotted owl (subspecies *S. o. caurina*) has been listed as “threatened” under the Endangered Species Act since 1990. Here we present a comprehensive look at genetic variation to elucidate the population histories of SO and invading western BO. Specifically, we present an improved SO genome assembly, based on 10x and Bionano Genomics data, along with 51 high-coverage whole-genome sequences including 11 SO from two subspecies (*caurina* and *occidentalis*), 25 BO, 2 confirmed and 13 potential hybrids. We identified potential hybrids based on intermediate morphology and found them to be a mixture of pure BO, F1 hybrids, and F1 x BO backcrosses. Unlike previous studies reporting asymmetries in the species-specific genders of the parents of F1 hybrids, we did not observe any significant asymmetry. Within species, we found that Western BO genetic variation is not simply a subset of the genetic variation in Eastern BO, suggesting that the two groups have been genetically isolated for longer (thousands of years) than previously suspected (80-130 years). Similarly, we found evidence of substantial genetic differentiation between the two SO subspecies. Finally, our analyses suggest that Northern SO experienced a moderate population bottleneck around the end of the last glaciation, while BO population sizes have always been large.

## Introduction

Spotted owls (SO, *Strix occidentalis*) occupy forests in western North America. There are three recognized, genetically distinct subspecies (Dawson et al. 1987; Fleischer et al. 2004; Barrowclough et al. 2005; Funk et al. 2008): the northern spotted owl (NSO, *S. o. caurina*), found from southern British Columbia southward to southern Marin County in California; the California spotted owl (CSO, *S. o. occidentalis*), found from approximately the Pitt River in northern California southward through the Sierra Nevada ranges to Baja, and northward along the coast ranges to San Francisco; and the Mexican spotted owl (MSO, *S.o. lucida*), found in mainland Mexico and the sky island forests of the south-western US deserts. Populations of all three subspecies have been declining for decades, leading the U.S. Fish and Wildlife Service to list the NSO and MSO as ‘threatened’ under the Endangered Species Act in the early 1990’s (Thomas et al. 1990). This act has led to changes in forest management practices across the Pacific Northwest, which have had an ongoing economic effect on the West Coast timber industry (Courtney et al. 2004). Although the listing of NSO was initially motivated by concerns over habitat loss (Forsman et al. 1984; Anderson and Burnham 1992), it is now clear that competition with the congeneric, invasive barred owl (BO, *S. varia*) poses an additional threat (Diller et al. 2016; Dugger et al. 2016). Observational data suggest that barred owls, previously inhabiting areas east of the Rocky Mountains and Great Plains, have expanded their range over the past 80-130 years (Dark and Gould, 1998; Livezey 2009a; Livezey 2009b) to include western North America, where they are sympatric with and out-compete spotted owls (Wiens et al. 2014). Barred owls continue to expand their range southward, currently overlapping with California spotted owls (CSO, *S. o. occidentalis*) as far south as Kern County, near Bakersfield, California.

Previous genetic work estimated an autosomal sequence divergence of 0.7% between spotted and barred owls (Hanna, et al. 2017). However, the two species have been shown to hybridize and backcross in the wild (Haig et al. 2004; Kelly and Forsman 2004; Hanna et al. 2018), leading to another concern for the conservation of spotted owls, that is, genetic invasion by barred owls. They hybridize primarily in areas where spotted owls greatly outnumber barred owls (Kelly and Forsman 2004). Observed interspecies mating pairs mainly involved a female BO with a male SO (Hamer and Forsman 1994; Haig et al. 2004; Kelly and Forsman 2004).

We had previously speculated that the unusual plumage pattern seen in some barred owls in their new western habitats was due to introgression with SO (see Figure 1, Hanna et al. 2018). However, analyses of low-coverage whole-genome sequence data from these birds suggest that the vast majority of these phenotypically unusual individuals were genetically purebred barred owls (Hanna et al. 2018). The question of how WBO evolved a unique plumage pattern in such a short period of time remains unresolved; one possibility is that barred owls in the western habitats (Western barred owls, WBO) may have diverged from barred owls in their original eastern habitats (Eastern barred owls, EBO) more than 130 years ago.

**Figure 1.**
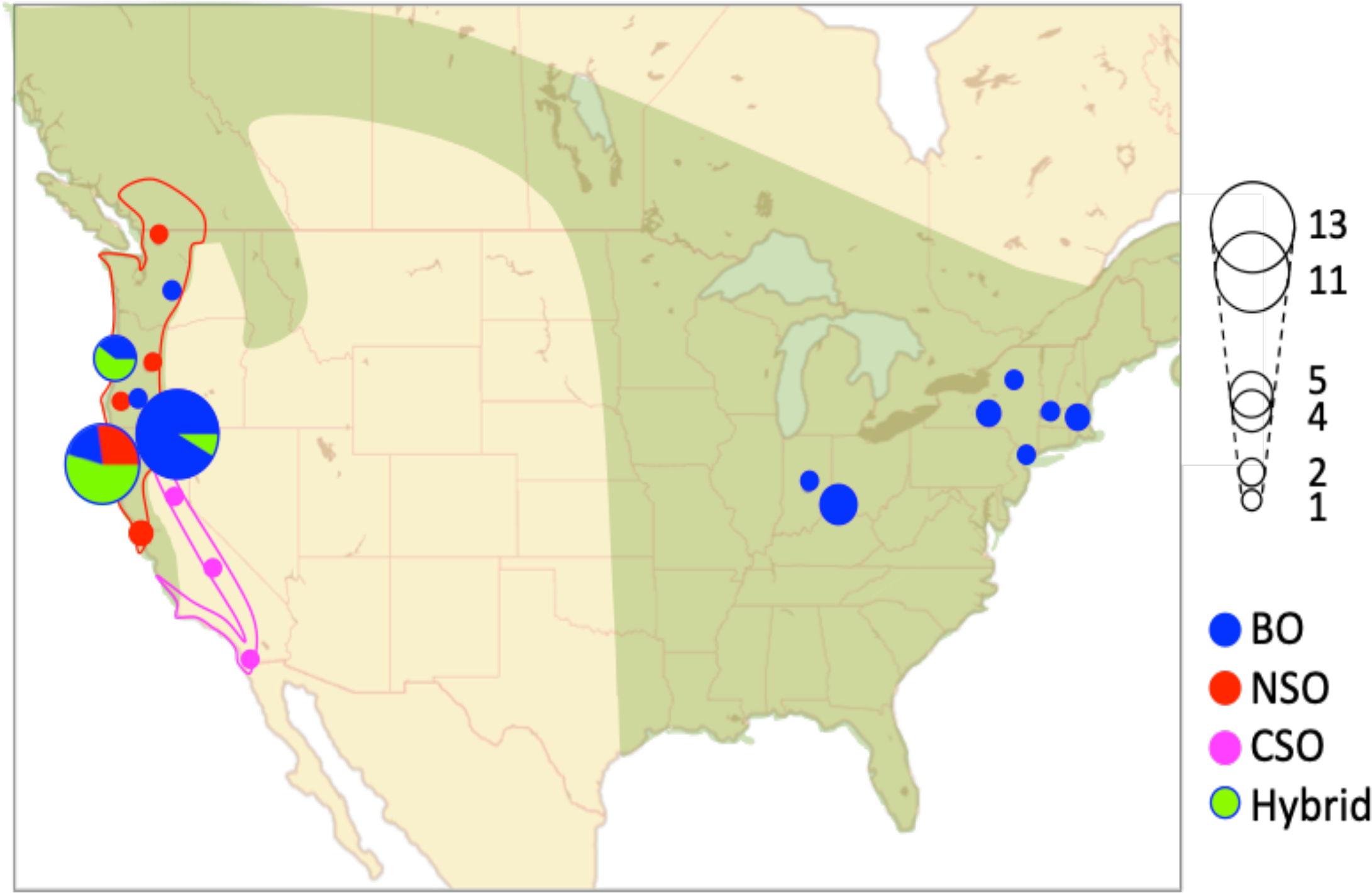
Geographic distribution of samples. Sampling locations of the 51 individuals in our study. Genetically identified northern spotted owls, California spotted owl, barred owls and hybrids are indicated by different colors. For locations with a high density of samples (e.g. Humboldt County and Siskiyou + Shasta County in California; Lane + Benton County in Oregon), the distribution of sampled individuals are visualized in pie charts. The size of circles and pie charts correspond to the number of samples. The range of barred owls was shown in green. The ranges for NSO and CSO are shown with red and magenta lines respectively.

In part to address this unresolved issue, we initiated a large-scale genomic study of spotted owl and barred owl genetic variation, incorporating an improved spotted owl genome assembly (using data from 10x Genomics and Bionano Genomics) and high-coverage whole-genome sequencing from 51 owls, including 8 NSO, 3 CSO, 12 EBO, 13 WBO, 2 known hybrids, and 13 owls of unknown ancestry. We use the resultant data to characterize patterns of genetic variation, divergence and hybridization both within and between owl species.

## Results

### New assembly of *S. occidentalis*

To facilitate high-resolution studies of population structure within and between *Strix* species, we improved upon the existing spotted owl genome, “StrOccCau_1.0”(Hanna et al. 2017), with10x Genomics (10xG) linked-read data and Bionano Genomics optical maps. For the new assembly, we used the same female *S. occidentalis* sample named Sequoia (hereafter simply Sequoia) that was used to construct the previous assembly(Hanna et al. 2017). Our new data resulted in a more contiguous assembly, “StrOccCau_2.0” (Table S1 and Figure S1), with the N50 scaffold size increasing from 4.0 Mb to 20.5 Mb (Assembly accession: XXXX).

### Description of the data

We generated high coverage (mean 31.70 X) whole-genome sequence data from 50 additional owl samples from various sampling locations (Figure 1, Table S2). For convenience, we used simple informal identifiers for these samples; the corresponding museum IDs are shown in Table S2. Since our new assembly still contains many small scaffolds, we used only the 97 scaffolds longer than 1 Mb for identification of autosomes. We also identified the sex of the samples using the CHD1 locus, a commonly used avian sex marker (Table S2). Among the 97 large scaffolds, 15 showed a read depth about half that of the other scaffolds in females (Figure S2). We identified these 15 scaffolds as partial Z chromosome sequences, and we classified the other 82 as autosomal (Table S3, S4). Because we did not find any W chromosome sequences in the set of scaffolds longer than 1Mb, we widened our approach to include scaffolds and contigs longer than 100 kb. We calculated the proportion of missing data for each scaffold and contig in males and females (Figure S3) and conservatively identified 44 putative W chromosome fragments in which the mean proportion of missing data in male individuals exceeds 99% (Table S5). The total lengths of scaffolds and contigs identified as autosomes, Z chromosome, and W chromosome are 1.09 Gb, 84.9 Mb, and 8.6 Mb, respectively. Detailed description of the autosomes and the Z and the W chromosomes are shown in Figure S4 and the Supplementary Materials (1.Genetic diversity on sex chromosomes). We restricted most of our analyses to the 82 large autosomal scaffolds. In these, we identified 17,385,299 biallelic SNPs, of which 8,543,351 had high-confidence genotype calls (GQ≥40) in all individuals, and 16,307,111 had a high-confidence genotype call in at least one individual.

### Characterization of samples

The range of morphological variation among hybrids and western barred owls often makes it difficult to distinguish them from each other based on appearance. Subspecies of SO have been historically recognized based on body size, plumage coloration and geographic range, but it is also not always clear (Haig et al. 2004; Barrowclough et al. 2005; Funk et al. 2008). Because of these reasons, we re-identified all the samples with genetic data using principal component analysis (PCA) on 870,053 autosomal SNPs without missing data after LD-pruning (Figure 2A). This analysis clearly revealed that SO, and eastern and western barred owl populations clustered separately, whereas hybrids are scattered between spotted owls and western barred owls (Figure 2A). PCA analysis also confirmed that eleven of our samples were spotted owls and twelve were eastern barred owls. Among the thirteen potential hybrids, four samples were clustered together with western barred owls, and nine were located between species clusters and confirmed as hybrids. In total, we confirmed seventeen western barred owl samples and eleven hybrids. Eight out of eleven hybrids were located in the middle on the x-axis, and the other three hybrids are scattered in positions closer to WBO. In the cluster of spotted owls, three individuals from southern California are slightly distanced from other samples, suggesting that they are California spotted owls. This distinction was further substantiated when we analyzed spotted owls separately from barred owls and hybrids (Figure 2B). Spotted owls formed two distinct groups, one consisting of the three samples from southern California, and the other consisting of eight samples from the northern coastal range, corresponding to the two subspecies of spotted owls, California spotted owls and northern spotted owls respectively (TableS2). A plot for barred owls only (Figure S5) showed that both of WBO and EBO were separated into small clusters, suggesting geographic population structure within each group.

**Figure 2.**
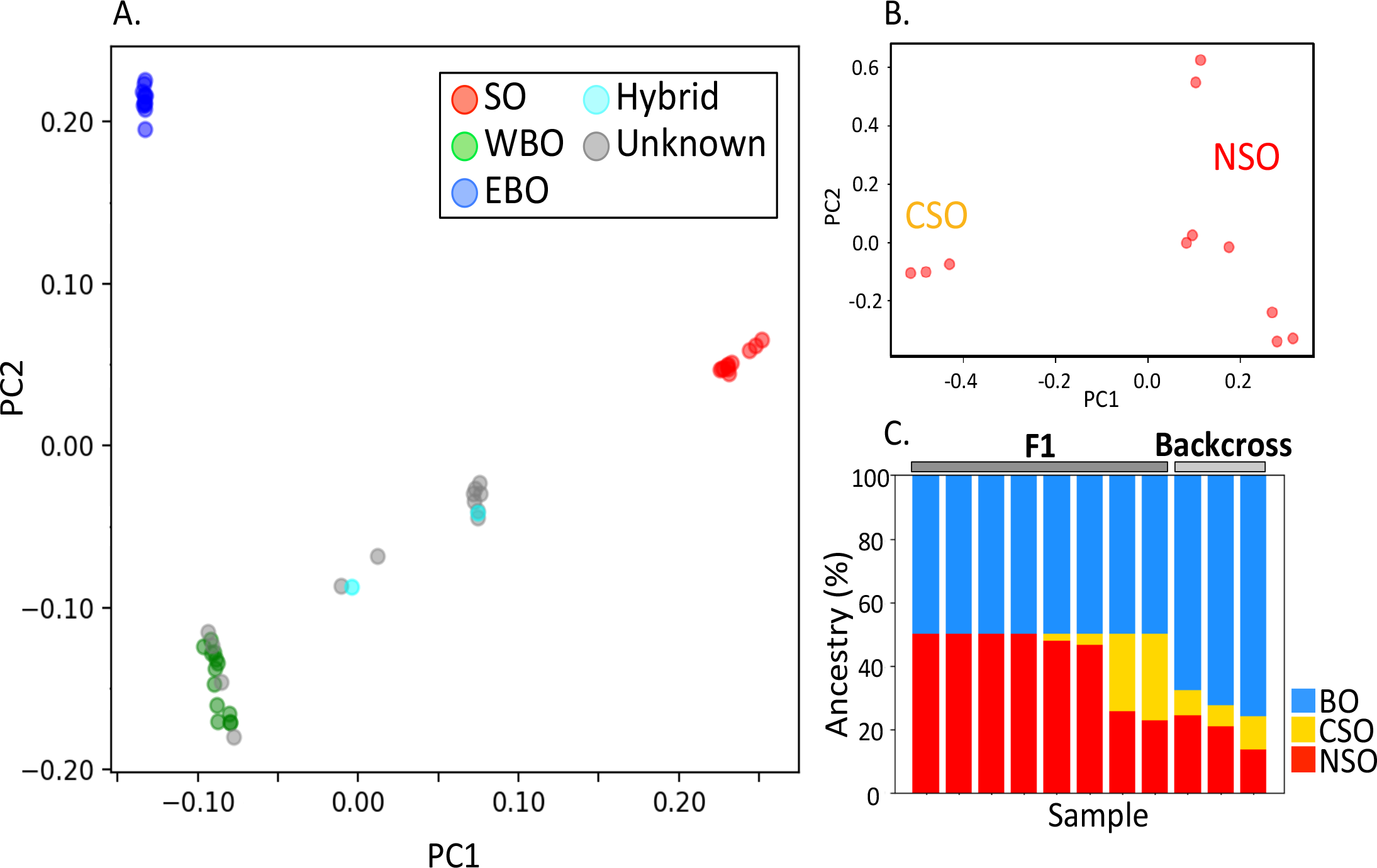
Principal component analysis (PCA). (A) PCA for 51 samples. Colors indicate the primary morphological identification of the samples. (B) PCA for 11 spotted owl samples. Clusters corresponding to the two subspecies; northern spotted owls (*S. o. caurina*) and California spotted owls (*S. o. occidentalis*) are shown. (C) Inferred ancestry of hybrids. Percentage of population-specific alleles is shown for each sample of hybrids.

To estimate the number of generations since hybridization, we tallied the number of genetic components from SO and BO for the 11 hybrid samples (Figure 2C, Table S6). We identified 2,484,025 apparent fixed differences between SO and BO and obtained percentages of spotted owl alleles for the hybrids. We then identified 772 apparent fixed differences between NSO and CSO at sites where no polymorphism had been observed in BO. We calculated the proportions of alleles coming from each of CSO and NSO in the spotted owl alleles. The eight samples, which were located in the middle of the x-axis on PCA plot (Figure 2A), showed 50.0% of spotted owl alleles, and all of the sites were heterozygous (Figure 2C, Table S6A), providing clear evidence of first-generation hybrids (F1). The other three samples showed 24.0 ∼32.3 % of spotted owl alleles, consistent with their closer positions to western barred owls on the PCA plot (Figure 2A). Because ancestry percentages can fluctuate in backcrossed hybrids due to recombination, the simplest explanation is that these three samples are F2 hybrids (backcrosses with BO), the expected proportion of which is around 25 % in spotted owl alleles. Between the two spotted owl subspecies ancestries, the genetic component from NSO was dominant over CSO and seen in all hybrids (Figure 2C and Table S6B), probably because the sampling locations of hybrids were in the range of NSO (Figure 1), although some are near the NSO/CSO hybrid zone.

### Diversity analysis

We sought to identify closely related individuals in order to avoid possible non-independence of close relatives or other effects of related individuals on our analyses of demography and genetic diversity (Materials and Methods, Supplementary Materials (2.Identification of close relatives)). As a result, we identified 8 parent-offspring pairs involving four different parents and eight offspring, one pair of full siblings, and one pair of closely related individuals, possibly half siblings (Table S7A). To avoid biasing our estimates of genetic diversity, we removed the four closely related samples ZRHG101, ZRHG123, ZRHG124 and ZRHG127 from our data for the diversity and demography analyses (Table S7B).

Using the 47 non-related individuals, we calculated genetic diversity for each population (Table S8). Consistent with a expected situation for threatened species, autosomal nucleotide diversities (π) of spotted owls were very small (1.27×10^-4^ for entire SO, 1.03×10^-4^ for NSO and 1.34×10^-4^ for CSO), while the nucleotide diversities of barred owls were more than 10 times higher (2.10×10^-3^ for entire BO, 1.94×10^-3^ for WBO and 2.14×10^-3^ for EBO). WBO, which is believed to have experienced a bottleneck during its recent invasion of the western US, showed slightly smaller π value than EBO. The nucleotide diversity between the two subspecies of spotted owls was 1.53×10^-4^, though the π between western and eastern populations of barred owls was 2.14×10^-3^, placing it in the same range as the π between spotted owls and barred owls, 6.02 x10^-3^. On the other hand, F_ST_ between northern spotted owls and California spotted owls was 0.253, while F_ST_ between eastern barred owls and western barred owls was 0.050, far smaller than the F_ST_ between the two species (0.765) (Table S9). Since both minor alleles and alleles with intermediate frequency can equally contribute to nucleotide diversity, π between populations reflects both the differentiation between the two populations and the population structure within each population. F_ST_ is commonly used for measuring differentiation between populations, though its estimator can be affected by asymmetry in sample sizes of the populations (Bhatia et al. 2013). In this case, the numbers of individuals of western and eastern barred owls are roughly equal (13 and 12 samples respectively), and F_ST_ should measure the differentiation between the two populations with accuracy. The large π between western and eastern barred owls detected here could be a result of their population structure.

The number of segregating sites is shown with their Tajima’s D values in Table S10. For spotted owls, Tajima’s D were negative (-0.47 ±1.06 for SO and -0.63 ±1.03 for NSO), suggesting that they experienced population expansions sometime ago, and that their decline in population size is too recent (starting ∼100 years ago) to be reflected to Tajima’s D as a positive value (Table S10). CSO showed a positive value of Tajima’s D, but we must note that the number of samples of them is small (n = 3) and they showed the highest variance. For barred owls, it is believed that they have kept a sufficiently large population size so far, and recent shrinkage of population size is not known. Consistent with this, Tajima’s D for the entire barred owl population and eastern barred owls were negative (-0.35±0.28 for BO and -0.52±0.25 for EBO). Western barred owls showed positive value (0.21±0.35), which is consistent with a founder event for WBO involving a small number of migrants from the eastern populations.

### Female Ancestry of hybrids

It has been suggested that hybridization between spotted owls and barred owls primarily involves male spotted owls pairing with female barred owls (Hamer and Forsman 1994; Haig et al. 2004; Kelly and Forsman 2004). One hybrid carrying a mitochondrial DNA haplotype of SO was previously reported (Haig et al. 2004), based on 524 bp of mitochondrial control region sequence, morphological traits, and vocalization. It was later found that both BO and SO have duplicated mitochondrial control regions (Hanna et al. 2017), thus establishing the need for higher resolution genetic methods in examining hybrids. To determine whether hybridization involving a female SO and a male BO happens with WGS data, we traced the maternal ancestry of the hybrids through the W chromosome. We identified 17,100 apparent fixed differences between SO and BO on the W chromosome, and we found that in two of the six female hybrids, all these sites were occupied by spotted owl alleles (Table S11), indicating that hybridization involving female spotted owls pairing with male barred owls also occurs. In the other four individuals, almost all sites were occupied by barred owl alleles. One of them (ZRH962) showed 0.12 % of spotted owl alleles, but this very low percentage suggests it is due to genotying error or incomplete assignments of alleles to species caused by small sample size.

### Inference of historical population size

We inferred historical changes of effective population size (N_e_) using SMC++ (Terhorst et al. 2017a) for NSO, WBO, and EBO (Fig 3). Because it is known to be difficult to infer very recent changes in N_e_, we focused on the last 200 - 500,000 generations (1000 – 2,500,000 years ago when generation time of 5 years is assumed). For NSO (Fig 3A), N_e_ gradually decreased to less than 10^3^ in the period from 50,000 -10,000 years ago, and then slowly recovered to ∼10^5^. The eight NSO trajectories in Fig 3A, which were each plotted using a different sample as a “distinguished” individual required in SMC++, are quite consistent with each other. Although we do not know actual generation time nor mutation rate of spotted owls, it is likely that the beginning of the recovery corresponds to the end of the last glacial period, as is frequently revealed in the studies of historical population size for temperate species (Nadachowska-Brzyska et al. 2015; Mays et al. 2018; Vijay et al. 2018). For barred owls, trajectories with different distinguished individuals showed greater variability (Fig 3B, 3C). EBO showed two types of curves, one with expansion around 50,000 years ago and the other with more constant population size (Figure 3B). These two patterns suggest two diverged populations of EBO. However, Weir and Cockerham’s F_ST_ (Weir and Cockerham 1984) between these two groups of “distinguished” individuals is extremely low (-0.0027), suggesting genetic diversity within the groups is higher than that between the groups. The members of the EBO groups with the alternate demographic patterns were mixed on the PCA plots (Fig2A and S5), and they don’t seem to correspond to the grouping based on the actual genetic components. The trajectories for WBO are similar to the constant pattern of EBO, but with declines in N_e_ at various times in the past (Figure 3C). It appears that populations of WBO split from EBO at various time points, but again, F_ST_ and π values between EBO and WBO are too small, 0.050 and 0.00214 respectively, to support such an ancient split. It is also inconsistent with our estimation of split time between EBO and WBO, as discussed further below. Although interpretation of the demographic trajectories for EBO and WBO are not clear (but see Supplementary Materials, 3. mtDNA analyses), they suggest that barred owls have successfully maintained large effective population size (e.g. N_e_ >10000) even during recent glacial cycles.

**Figure 3.**
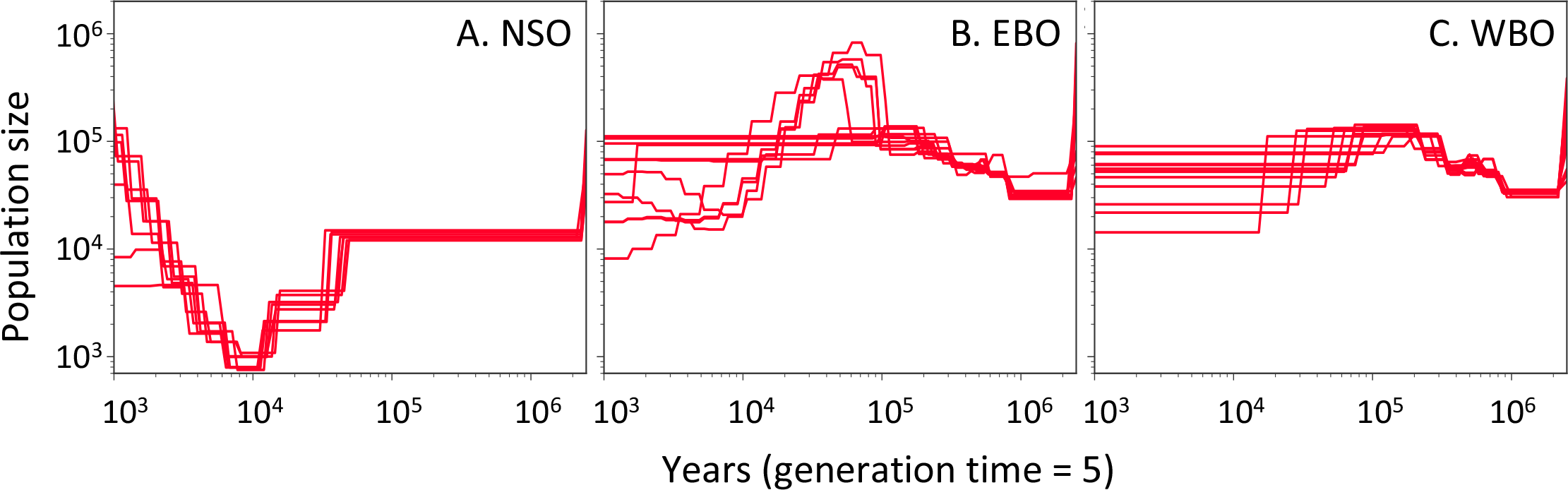
Demographic history inferred by SMC++ for (A) northern spotted owls, (B) eastern barred owls, and (C) western barred owls. Each trajectory was drawn with different distinguished sample(Terhorst et al. 2017a). A mutation rate of 4.6 * 10^-9^ / bp per generation and a generation time of 5 years were used.

### Split time between populations

Given the likely founder events associated with the range expansion of barred owls into western North America, it is unclear whether the within-species differentiation between EBO and WBO seen with PCA and F_ST_ reflects older divergence between groups or simply a recent bottleneck. The timing for the WBO population to have started to migrate to western North America is believed to be between 80 and 130 years ago (Livezey 2009a). Several methods such as SMC++ (Terhorst et al. 2017a), ∂a∂i (Gutenkunst et al. 2009) and PSMC (Li and Durbin 2011), are used to estimate split times of populations, but none of them are able to estimate split times over such recent history. Our quantitative approach to addressing this issue focuses on private variants, which are likely to be more informative than common variants about recent demographic changes. For all possible groups involving all of the individuals of a source or “focal” population (e.g. EBO) and a single individual from a derived or “test” population (e.g. WBO), we tabulated the number of alleles present in each individual but not segregating in the remaining individuals. For example, we assumed all possible groupings of 12 EBO and 1 WBO; for each group, we counted the number of alleles present in each individual but not segregating in the remaining 12, that is, we counted singletons and private homozygotes. This asymmetrical sampling scheme is useful in reducing the effects of population history (e.g., population bottlenecks) in the test population. The number of private alleles in a test individual reflects the length of time since the population split, and the averaged number of private alleles for a focal individual represents the depth of the genealogy within a focal population. If WBO had been isolated from EBO for a substantial length of time, we would expect the single WBO sample to contain more private variants than the EBO samples. This is exactly what we observed. We then took the ratio of the average number of private alleles in a test individual to the average number of private alleles in a focal individual and compared this with the expected ratios for certain hypothesized split times derived from coalescent simulations (see Materials and Methods for further detail). Using singletons is potentially challenging because they are also candidates for sequencing errors. However, the effects of sequencing errors is likely to be minor because of the high coverage of the data.

The numbers of private alleles in the EBO samples range from 129,000 – 145,000 (with a mean of 133,223 ±238 across samples and groups), while the corresponding number of private alleles in a single WBO test sample range from 149,000 – 161,000 (mean 152,383 ±2942) (Figure 4A, Table S12A), yielding an observed ratio of 1.14. Assuming a simple split model, we estimate the time of divergence between EBO and WBO as 0.0029 *4N_e_ generations (Figure 4C). Based on an assumed generation time of 5 years (see Materials and Methods, Generation time for analyses), this divergence time is estimated to have occurred around 7000 years ago. Even when allowing for uncertainty in our generation time estimate, our results appear to be at odds with the commonly assumed scenario of a very recent divergence (i.e., within the past 80-130 years) of WBO from the EBO population. However, using these methods, we are unable to distinguish between scenarios in which WBO are very recent descendants of an unsampled group from eastern North America and scenarios in which barred owls actually colonized some portion of western North America earlier than suggested by the historical record.

**Figure 4.**
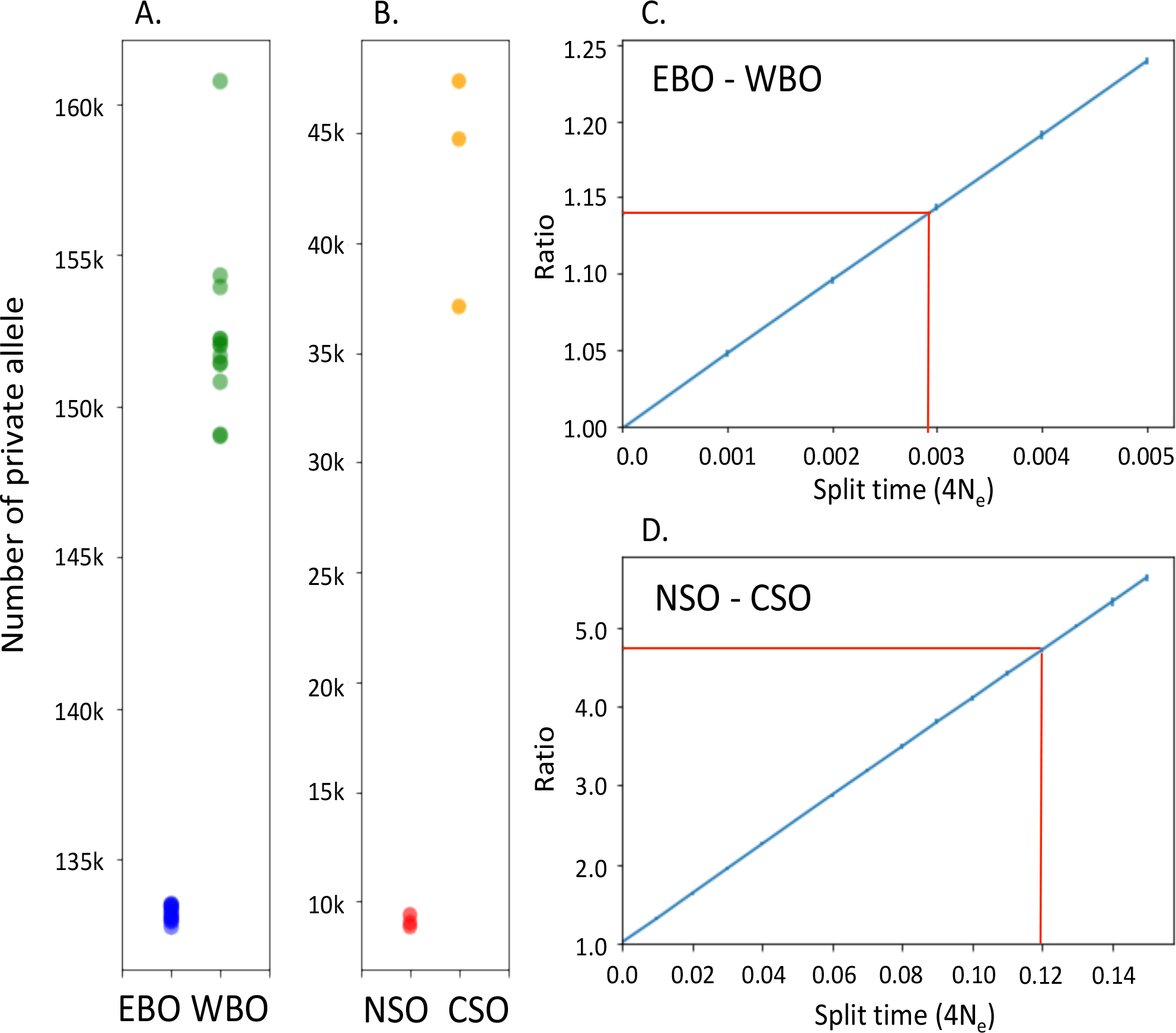
Estimation of split time between populations. (A) Observed numbers of private alleles of WBO and EBO were compared in 13 groups. A green dot shows the number of private alleles in each WBO sample, whereas a blue dot indicates the averaged number across the 12 EBO samples in a group. (B) Observed numbers of private alleles in CSO samples compared with average numbers of private alleles in NSO in 3 groups. (C) (D) Simulated (expected) ratio of number of private alleles in a test individual relative to the focal population is plotted against split times between populations. Red lines indicate the observed ratio for BO (C) and SO (D) of 1.14 and 4.74, respectively.

Using the same approach, we find a larger difference in private alleles between NSO (mean 9,082 ±271) and CSO (mean 43,068 ±5334) (Figure 4B, Table S12B). These data correspond to a divergence time of 0.12*4N_e_ generations (Figure 4D), or around 14,000 years ago under the assumption of generation time of 5 years. This estimate is consistent with the possibility that NSO and CSO differentiation was driven by occupation of different forest refugia during the last ice age. Since there has been some genetic migration between CSO and NSO in recent years, we expect CSO private allele counts to be positively correlated with distance from the NSO home range. This is exactly what we observed, with the closest CSO sample (ZRHG103 from Nevada County) having the smallest CSO private allele count (37,100) and the farthest CSO sample (ZRHG104 from San Diego County) having the largest CSO private allele count (47,372; Table S12).

## Discussion

Owls of the genus *Strix* have long been of great interest to many groups, partly because they are large, charismatic vertebrates, and partly because of the ecological, environmental and economic consequences of listing NSO under the Endangered Species Act. While there have been several genetic studies of spotted owls over the past 20 years (Barrowclough 1990; Barrowclough et al. 1999; Haig et al. 2001; Haig et al. 2004; Barrowclough et al. 2005; Barrowclough et al. 2011), recent advances in molecular biology and computational genetics now enable us to generate and analyze the complete genome sequences of spotted owls, barred owls, and their hybrids. Our analysis of 51 individuals is the largest genomic study of high-coverage genomes in spotted owls, barred owls, and hybrids, and it allows us to elucidate at high resolution the population dynamics both within and between species.

Though we had a limited number of spotted owl samples, multiple analyses confirmed that CSO and NSO are well-defined evolutionary groups, with each subspecies containing a substantial amount of private genetic variation. These data derived from analyses of whole-genome sequences provide additional evidence in support of management strategies that consider each spotted owl subspecies as a separate evolutionarily significant unit. Our divergence time estimate (14,000 years ago) is roughly contemporaneous with the time of the last glacial maximum, during which time suitable forest habitat may have been somewhat fragmented.

More unexpectedly, we found substantial differentiation between WBO and EBO that is inconsistent with a separation time of between 80 and 130 years ago. There are two plausible explanations for this observation: First, we do not know *where* this divergence may have occurred. Given our limited sampling of EBO individuals, it is possible that there is a substantial amount of genetic variability within EBO, and that there exists an unsampled EBO population that is directly ancestral to extant WBO individuals. Work by Barrowclough et al. (2011) using mitochondrial data (Barrowclough et al. 2011) suggests that there is substantial variation within EBO. Although we confirmed that our samples included the population structure observed with mtDNA, it is still possible that our data do not fully cover the range of EBO diversity (Supplementary Materials, 3. mtDNA analyses). Second, it is possible that the observational data (i.e., the first observed sightings of barred owls in different western North America locales) are inaccurate or incomplete. Regardless, our data clearly refute a scenario in which the WBO samples are very recently derived (i.e., within the past 130 years) from a panmictic population of EBO (as encapsulated by the 12 EBO samples examined in this study). Since our analyses focus on barred owl-specific variants, we believe that this ambiguity in barred owl population history can be explained only by distinct evolutionary histories of the sampled WBO and EBO individuals. Finally, we would like to emphasize that the results of our methodology are insensitive to unknown facets of WBO population history, such as any potential population bottleneck associated with the founding of WBO populations.

Our use of whole-genome sequencing also allowed us to classify potential spotted vs. barred owl hybrid individuals. In contrast to a general lack of hybridization between NSO and WBO across much of their range (Hanna et al. 2018), hybridization appears to be a more significant phenomenon at the leading edge of the WBO expansion into regions such as the Northern Sierras where WBO are still rare (Kelly and Forsman 2004), yet we have little understanding of the overall fitness and ultimate fates of hybrid individuals. Out of 15 potential hybrids in our sample, we identified four WBO, eight F1 hybrids and three F1 x barred owl backcross individuals. This distribution confirms our previous findings that phenotypically distinct barred owls from California are not necessarily hybrids (Hanna et al. 2018). However, in a departure from previous studies (Hamer and Forsman 1994; Kelly and Forsman 2004), we found that male SO x female BO and male BO x female SO offspring occur at roughly equal frequencies. We also observed that the spotted owl contribution to these hybrid individuals included NSO and F1 CSO x NSO individuals. Our results, however, cannot directly address the absence of later-generation hybrids between spotted and barred owls. It is unclear at this time whether these later-generation hybrids are not found due to hybrid incompatibilities, or whether further sampling of potentially hybrid individuals would uncover a deeper collection of multi-generation hybrids. Additional in-depth studies of potential spotted vs. barred owl hybrids will be necessary to answer this question.

## Materials and Methods

### Assembly of the new reference genome

To obtain an improved spotted owl reference genome, we generated a hybrid (10x Genomics and Bionano Genomics) assembly following the approach in Levy-Sakin et al (Levy-Sakin et al. 2019). Briefly, we obtained high-molecular-weight DNA from blood sample of Sequoia, and used this to generate a 10x Genomics (10xG) linked-read library (using their Chromium system) and Bionano genome maps (using their Irys system). Instead of generating a single genome map with the enzyme Nt.BspQI, we generated two sets of Bionano genome maps with the enzymes Nt.BspQI (New England Biolabs (NEB), Ipswich, MA, USA) and Nt.BbvCI (New England Biolabs (NEB), Ipswich, MA, USA). The 10xG library was sequenced to an average depth of ∼60X and assembled using Supernova v1.1(Weisenfeld et al. 2017). We then generated hybrid scaffolds using the Bionano genome maps to bridge Supernova scaffolds (see Levy-Sakin et al. 2019 for further details).

### Sequence data

We utilized whole-genome sequencing data from a previous study (Hanna et al. 2017) for Sequoia (Table S2). For the other fifty samples from various sampling locations (Figure 1), we extracted genomic DNA following the method described in Hanna et al. (2017), prepared whole genome libraries using a Nextera DNA Sample Preparation Kit (Illumina) and obtained high-coverage paired-end sequences from MedGenome, Inc. using a mix of different Illumina HiSeq machines. For convenience, we used simple informal IDs for all the samples, though corresponding museum IDs are shown in Table S2. The location map was made with leaflet package (Graul 2016).

### Alignment and processing of data

We processed the paired-end data from the whole genome libraries of the fifty-one samples. We used Picard 2.19.0-SNAPSHOT in Genome Analysis Tool Kit (GATK) version 4.1.2.0 (McKenna et al. 2010; Depristo et al. 2011; Van der Auwera et al. 2013; Poplin et al. 2017) to remove adapter sequences. Then we modified the pipeline, processing-for-variant-discovery-gatk4.wdl supplied by the GATK as a Best Practice of GATK4, to use in our local environment. We aligned the trimmed paired reads to our new reference “StrOccCau_2.0_nuc_finalMito.fa” using bwa mem version 0.7.12-r1039 (Li 2013). We performed two rounds of base quality score recalibration (BQSR) in GATK4 using SNPs previously identified by Hanna et al. (2017) (Hanna et al. 2017).

### Variant calling and filtering

We called variants using the GATK4 HaplotypeCaller for each of the fifty-one samples, and then performed joint genotype calling with the GATK4 GenotypeGVCFs tool for all samples included as simultaneous inputs. We used the GATK4 VariantFiltration to remove variants more extreme than a p-value of 3.4e-6 in Hardy-Weinberg equilibrium, which phred-scaled is 54.69.

We followed the guidelines of GATK for hard filtering (https://software.broadinstitute.org/gatk/documentation/article?id=23216#2, https://software.broadinstitute.org/gatk/documentation/article?id=11069) to retain only high-quality, biallelic single nucleotide polymorphisms (SNPs). First, we used the GATK SelectVariants tool to extract the single nucleotide polymorphisms (SNPs) from the raw VCF file. Then we filtered the SNPs using the GATK VariantFiltration tool with options ‘--filterExpression “QD < 2.0 || FS > 60.0 || MQ < 40.0 || MQRankSum < -12.5 || ReadPosRankSum < -8.0 || SOR > 3.0”’. Then we removed any variants that fell within repetitive or low complexity regions using BEDTools version 2.25.0 (Quinlan and Hall 2010). To retain only biallelic sites, and to remove variants on the mitochondrial genome, we used the GATK SelectVariants tool with the “--restrict-alleles-to BIALLELIC -XL Sequoia_complete_mtGenome --exclude-filtered” options. We calculated the mean and standard deviation of the total unfiltered read depth across all samples per site, and removed all the variants exceeding the mean coverage plus five times the standard deviation, as suggested by the GATK documentation. In addition to these basic filters, we filtered out individual variants with the minimum quality of assigned genotype (GQ) smaller than 40.

We also removed the sites with missing data for all the analyses below except for the diversity analysis.

### Sex identification

A previous study (Hanna et al. 2017) identified scaffolds 806 and 4429 on their reference genome “StrOccCau_1.0_nuc.fa” as the scaffolds including matched sequences with CHD1Z or CHD1W, which are known as markers of sex for avian species (Fridolfsson and Ellegren 1999), suggesting that scaffolds 806 and 4429 are sequences from the Z and W chromosomes respectively. We identified a corresponding scaffold for each of them in our reference genome “StrOccCau_2.0_nuc_finalMito.fa” with NCBI BLAST and checked CHD1Z and CHD1W sequence were there. Using the difference in read depth on the correspondents, we identified sex for each of the fifty-one samples.

### Autosome and sex chromosome identification

Birds have the ZW sex-determination system, where the female is the heteromorphic sex (ZW) and the male homomorphic (ZZ). Since our reference genome is female, reads from both of the sex chromosomes were mapped to it. For identification of the Z chromosome and autosomes, we calculated the mean read depth for each scaffold in each sample. Then we took the averaged read depth of each scaffold across samples for males and females. Based on the assumption that the read depth of the Z chromosome would be half in females as in males, we searched for scaffolds with approximately half the averaged read depth across variants in female samples as in male samples, and identified them as sequences that likely map to the Z chromosome. We also identified the scaffolds with similar read depth in males and females as autosomes.

For identification of the W chromosome, we quantified the amount of missing data, because in males the variants on the W chromosome should be missing. To exclude low-quality regions, we applied a GQ filter of ≥ 40 (using vcftools (Danecek et al. 2011)) and removed variants where more than half of the samples had missing genotypes. (Note that exactly half of our samples are female.) For the final set of variants, we calculated percentages of missing data for each scaffold and contig of each sample. We searched for scaffolds or contigs where more than 99% of sites are missing in all male individuals in the pool of scaffolds and contigs longer than 100 kb, identifying them as W chromosome sequences.

### Principal Component Analyses (PCA)

For PCA analysis, we used only autosomal scaffolds. We also pruned variants to leave variants with minor allele frequency at least 1 %, with no pairs remaining with r^2^ > 0.2 for the sets of samples, using PLINK (Purcell et al. 2007). Then we performed PCA with PLINK.

### Identification of close relatives

We sought to identify closely related individuals in order to avoid possible non-independence of close relatives or other effects of related individuals on our analyses of demography and genetic diversity. Since we do not have phased haplotypes for the sequenced genomes, we could not use standard Identity-By-Descent (IBD) methods for detecting close relative pairs. Instead, we calculated the kinship coefficient (phi) (Manichaikul et al. 2010) and proportion of the sites where two individuals share zero alleles identical by descent (proportion of zero IBS) for each pair of individuals within and between populations (see Supplementary Materials for further detail).

### Diversity analyses for autosomes

For diversity analyses, we removed the 4 samples, which have closely related samples within species as described in Supplementary Materials (Table S7B). We also removed all the variants from any individual with the GQ score smaller than 40 with vcflib (Garrison E). Using variants on autosomes, we calculated the number of segregating sites, and Tajima’s D (Tajima 1983) for each population with vcftools (Danecek et al. 2011). We measured Weir and Cockerham’s *F_ST_* with PLINK (Weir and Cockerham 1984; Purcell et al. 2007), and calculated nucleotide diversity within and between populations or groups using our python scripts. We also calculated Hudson’s F_ST_ (Hudson and Slatkint 1992) to compare with those for the sex chromosomes using our python scripts (Supplementary Materials, 1.Genetic diversity on sex chromosomes).

### Ancestry analyses

To calculate a percentage of spotted owl ancestry in hybrids, we identified apparent fixed differences between spotted owls and barred owls in our samples. For each known or potential hybrid, we calculated the mean percentage of ‘spotted owl alleles’ at these fixed differences as well as the mean heterozygosity.

Similarly, we identified apparent fixed differences between NSO and CSO, at sites where no polymorphism is observed in barred owl samples, to estimate the percentages of sub-specific spotted owl ancestries in hybrids. Assuming one of the parents of each hybrid is a barred owl, we tabulated the mean percentages of ‘NSO alleles’ across these NSO vs. CSO fixed differences for each hybrid individual.

To estimate the percentage of spotted owl ancestry on the W chromosome in female hybrids, we extracted scaffolds and contigs identified as partial W chromosomal sequences from females, using the filtered vcf file of the W chromosome described above. Then we examined the fixed differences between spotted owls and barred owls there. We then calculated the proportion of ‘spotted owl alleles’ across these fixed differences for each female hybrid.

### Generation time for analyses

For generation time of spotted owls, multiple estimations have been made so far, including two (Gutiérrez and Franklin 1995), five (Barrowclough and Coats 1985; Barrowclough et al. 1999), or ten years (USDA Forest Service 1992; Noon and Biles 1990). When we considered the reported low rate of successful breeding in the early stage of spotted owls (Forsman et al. 2002), 5 - 10 years seem to be proper. We used a mean generation time of 5 years to be conservative to scale the split time estimation and to scale the estimation of population size history.

### Inference of population size history

To estimate population size and infer demographic histories for northern spotted owls, eastern barred owls and western barred owls, we used SMC++ version 1.15.2 (Terhorst et al. 2017b). We applied the mutation rate of collared flycatcher (*Ficedula albicollis*), 4.6E-9 per site per generation (Smeds et al. 2016) and a generation time of 5 years (see above). Because it is known to be difficult to infer very recent changes in N_e_, we focused on the last 200 - 500, 000 generations (1000 – 2,500,000 years ago when generation time of 5 years) is assumed for NSO, WBO, and EBO. We estimated the population size history multiple times using every different sample in a population as a distinguished individual. We also calculated it for EBO and WBO in the same way, though we couldn’t do it for CSO because of its small sample size.

### Inference of split time of populations

To infer the split time between WBO and EBO, we used alleles private to an individual. We defined a source population and a population derived from a source population as “focal” and “test” populations. We made a group of individuals consisted of all of the individuals of source or “focal” population (e.g. EBO) and single individual from derived or “test” population (e.g. WBO). We used this asymmetrical sample groups to reduce the effects of demographic facets within the test population, such as a population bottleneck. For all possible groups, singletons and private homozygotes (i.e., when one individual is homozygous for one allele and all other individuals are homozygous for the other allele) in each individual are counted. All singletons were counted once, while private homozygotes were counted twice. We count private homozygotes as well as singletons to make it robust against inbreeding since the number of singletons is quite vulnerable from inbreeding. For each group, we recorded the number of private alleles for a test individual and the mean number of private alleles across focal individuals. Here, the number of private alleles in a test individual reflects the length of time since the split of the two populations, while the averaged number of private alleles for a focal individual represents depth of the genealogy within a focal population. If the split of the two populations occurred very recently, the variation in a test individual will be only a subset of that in a focal population, leading the two numbers very close. However, if a test population has been isolated from a focal population for a substantial length of time, we would expect a single test individual to contain more private alleles than a focal individual. Then the mean numbers of private alleles for a test and a focal individual were calculated across all the possible groups, to take a ratio of the averaged number for a test individual to that of a focal individual. We compared the observed ratio to the expected ratio obtained from simulations (described below) to estimate the split time between focal and test populations.

In this case, using the 12 EBO samples and the13 WBO samples, we took all the 13 possible groups of a single WBO sample (a test individual) and 12 EBO samples (focal individuals), and tabulated the number of private alleles (from the filtered biallelic set of high-confidence biallelic SNPs (GQ>=40) without missing data described above) for each sample in each group. For the EBO samples, we averaged the number of private alleles across 12 individuals. Then we averaged the numbers across all the 13 groups for EBO and WBO, and took the ratio of the value for WBO to the value for EBO. An analogous approach was used to derive the ratio across the 3 possible sets of one of the three CSO (as a test population) samples and 8 NSO (as focal population) samples.

To obtain the expected values of the ratio, we simulated a simple even split of two populations with constant population size under standard neutral model, using a standard coalescent simulator (ms (Hudson 2002)). With an option of “./ms 50 100000 -t 22.1 –r 22.1 10000 -I 2 24 26 -n 2 1.0 -ej <split time> 2 1” for BO, we simulated 100,000 short segments of 10 kb to mimic a entire genome of 1 Gb, given a mutation rate of 4.6 x 10^-9^ / bp per generation (Smeds et al. 2016), the same magnitude of recombination rate and a effective population size of 120000. For SO, we used the command line of “./ms 22 10000 -t 11.0 –r 11.0 100000 -I 2 16 6 -n 2 1.0 -ej <split time> 2 1” to simulate 10,000 sequences of 100 kb, with an effective population size of 6000. We counted private alleles in a group of one test individual and a focal population in exactly the same way we did for the observed data. We calculated the mean number for a test and a focal individual across the groups and took the ratio of the numbers of private alleles for a test individual to that for a focal individual. We took the ratio as a function of the split time T (parameterized in units of 4N_e_ generations where N_e_ is the effective population size), across increments of 0.001 (for Barred Owl comparisons) or 0.01 (for Spotted Owl comparisons). We took 100 replications of the simulation for each of BO and SO. We then used a method of moments approach (with linear interpolation) to estimate split time from the observed ratio of test vs. focal individual private alleles.

To convert coalescent time units into years, we assumed generation time of 5 years (see above), a mutation rate of 4.6 x 10^-9^ / bp per generation (Smeds et al. 2016), and the effective population sizes of 120000 and 6000 for EBO and NSO respectively, estimated from the nucleotide diversity of these populations (Table S8).

## Supplementary materials

The supplementary text and supplementary tables and figures are found in Supplementary Materials except for Table S2∼S5 and S12, which are attached as excel files.

## Data availability

Raw sequence reads are available from the NCBI Sequence Read Archive (SRA) run accessions XXXXX. The assembly, “StrOccCau_2.0”, and the vcf file used in this study are available at XXX.

## Acknowledgement

This work was supported by a University of California President’s Research Catalyst Award and an unrestricted gift (to JDW) from Sierra Pacific Industries.

Map data copyrighted OpenStreetMap contributors and available from https://www.openstreetmap.org.

