## Supplementary information for "Genomic variation and population histories of spotted (*Strix occidentalis*) and barred (*S. varia*) owls"

#### Supplementary Materials

1. Genetic diversity on sex chromosomes
2. Identification of close relatives
3. mtDNA analyses
    3-1. Extraction of mitochondrial variants
    3-2. Phylogenetic analysis of the mitochondrial non-coding region
    3-3. Identification of mitochondrial lineages

Captions for Supplementary Figures
Captions for Supplementary Tables
Supplementary reference
Supplementary figures
Supplementary tables

##### 1. Genetic diversity on sex chromosomes

For the purpose of confirming the consistency in the identification of autosomal and sex chromosomal variants, we performed diversity analyses on sex chromosomes as well as on autosomes. As in diversity analyses on autosomes, we removed the 4 samples, which have closely related samples within species (Table S7B). We also removed all the variants from any individual with the GQ score smaller than 40 with vcflib (Garrison E). We calculated π within and between populations (Figure S6, TableS13) with PLINK (Purcell et al. 2007), and measured Hudson's $F_{ST}$ (Hudson and Slatkint 1992) with our python scripts.

The average levels of nucleotide diversity within species for the Z chromosomal variants were significantly lower than autosomal variants, whereas that for the W chromosome was very low (Figure S6A). Since the effective population size for the Z chromosome and the W chromosome are three-quarters and one-quarter of autosomes respectively, the neutral equilibrium expectation of relative genetic diversity of Z and W chromosome to autosomes are also 0.75 and 0.25. Here, the relative genetic diversity of the Z chromosome to autosomes is approximately 0.4 for spotted owls (0.395 for SO, 0.427 for NSO, and 0.367 for CSO) and 0.5 for barred owls (0.525 for BO, 0.529 for WBO, and 0.522 for EBO). Deviation from this expectation can be caused by many factors, such as different mutation rate between sexes, sex-biased demographic events, and genetic drift (reviewed in Sayres 2018). Another possible factor is a recent decline in population size. Because of its smaller effective population size, a decrease in population size reduces diversity more on the Z chromosomes than on autosomes (Pool and Nielsen 2007). Purifying selection against recessive deleterious mutations is also a factor reducing the diversity across the Z chromosomes, because they are always exposed in female individuals. A combination of these factors would be the truth.

The observed nucleotide diversity on the W chromosome is extremely low for both of spotted owls and barred owls (relative diversity to autosomes is 0.047 ~ 0.072 for spotted owls and 0.010 ~ 0.017 for barred owls) (TableS8, TableS13B), compared with the equilibrium neutral expectation for the W chromosome of 0.25. Extremely low W-linked variation was also reported for other avian species (for flycatchers (Smeds et al. 2015) and for chicken (Berlin and Ellegren 2004; Moghadam et al. 2012). The reduction of diversity was also observed for the human Y chromosome (5 -10 lower diversity than autosomes), which has been explained by selective sweeps on testis-specific genes (Wilson Sayres et al. 2014). Selection is also a likely explanation for low diversity on the avian W chromosome. Recombination does not occur in W-specific or Y-specific regions, selective sweep for loci can affect the entire chromosomes.

We also observed higher diversity between species for the Z chromosome than that for autosomes (Figure S6B, Table S13A). This is a commonly observed pattern and supported theoretically as "fast-Z effect" caused by faster fixation of recessive beneficial alleles on a hemizygous chromosome in one sex. $F_{ST}$ between species showed that alleles on the W chromosome are almost fully sorted with a low proportion of shared polymorphism (Figure S6C), and that $F_{ST}$ of the Z chromosome is intermediate to autosomes and W chromosome. This is consistent with the expected small size of $N_e$ for the W and Z chromosome. When the effective population size is smaller, the rate of lineage sorting is faster. In total, our observation on genetic diversity of the sex chromosomes is quite consistent with the previous observations on avian sex chromosomes, suggesting the sorting of the autosomal and sex chromosomal variants in our data was reliable.

**2. Identification of close relatives**

Since we do not have phased haplotypes for the sequenced genomes, we could not use standard Identity-By-Descent (IBD) methods for detecting close relative pairs. Instead, we used Identity-By-State (IBS) calculations as follows. We calculated kinship coefficient (phi) (Manichaikul et al. 2010) and proportion of the sites where two individuals share zero alleles (proportion of zero IBS sites, IBS0) for each pair of samples within and between populations. We used the proportion of zero IBS sites to distinguish parent-offspring pairs and pairs of full siblings, since the expected phi values of these two categories are the same, 0.25. The expectation of IBS0 for parent-offspring pairs is always 0, while that of full siblings cannot be 0. Combining phi and IBS0, we detected closely related samples (within 2$^{nd}$ degree relatives) and estimated their relationship.

We used the “relatedness2” option in vcftools(Danecek et al. 2011) to calculate statistics related to phi and to count the total number of sites where no alleles are shared between the two individuals of a pair. Using the statistics from vcftools, we calculated an estimator of phi, with Equation (11) in Manichaikul et al. 2010(Manichaikul et al. 2010). Originally they use Equation (9) for within-family relationship checking and Equation (11) for between-family relationship checking. Because Hardy-Weinberg Equilibrium (HWE) among SNPs is assumed for Equation (9), Equation (11) was derived in order to guard against potential estimation inflation due to departure from individual-level HWE. If there is no departure from individual-level HWE, the estimator of Equation (9) is no larger than that of Equation (11). We used Equation (11) for our data, because we observed departure from HWE as a distance from the diagonal line in SO and hybrids (Fig S7C -E), while in BO the two estimators are almost the same, indicating BO are ideally under individual level of HWE (Fig S7A and

S6B).

Their major concern in Manichaikul et al. 2010 was that the violation of HWE in the direction of too little homozygosity (due to reasons such as genotyping errors, recent admixture in a mixed population, or removing Mendelian errors in families) makes the estimator (Equation 9) over-estimate phi. To correct the effect of violated HWE, they used the smaller (the better) of the observed heterozygosity rates of a pair of samples (the number of heterozygous sites in a individual divided by the total number of non-missing markers for the pair of individuals) as an alternative to as expected heterozygosity for Equation (11).

In our case, the major concerns of violation of HWE are the extremely low genetic diversity of SO predicted from their current endangered status and the decreased homozygosity in hybrids due to the recent admixture. To mitigate these effects of violated HWE, we used the larger of the observed heterozygosity rates of a pair involving a SO sample as an alternative to expected heterozygosity in Equation (11) and used the smaller for a pair that doesn't involve SO. For the pairs involving a SO and a hybrid samples, phi values are potentially inflated, since the too high heterozygosity of hybrids would be used as the larger heterozygosity in a pair. To examine this possible inflation of phi, we used the number of segregating sites and the number of zero IBS sites.

First, we searched for closely related samples within 2$^{nd}$ degree relatives in the pairs of individuals within NSO (Fig S8A), CSO (Fig S8B), EBO (Fig S8C), and WBO (Fig S8D). No related individuals were detected from the kinship coefficient for NSO, CSO, and EBO. The absence of parent-offspring pairs in these populations was also confirmed the fact that all the IBS0 values of these populations are significantly larger than 0. For the pairs in WBO (Fig S8D), six pairs of parent-offspring or siblings were identified by phi. Among the six pairs, four showed almost 0 of IBS0 values ($5.73 \times 10^{-6}$ – $1.75 \times 10^{-5}$), while two showed significantly larger values ($2.76 \times 10^{-2}$ and $4.02 \times 10^{-2}$). We concluded that the former are parent-offspring pairs and the latter are siblings. The small phi values of the four pairs would correspond to genotyping errors. For the two pairs of siblings, we identified the pair of ZRHG114 and ZRHG123 (phi = 0.248) as full siblings and the pair of ZRHG126 and ZRHG127 (phi = 0.127) as close relatives, likely half-siblings, since the expected phi value for full siblings is 0.25 and that for half-siblings is 0.125.

Next, we searched for related individuals in the pairs between BO and hybrids and pairs between SO and hybrids. For the pairs between BO and hybrids, no close relatives were found (Fig S9A and S9B), while for the pairs between SO and hybrids, a cluster of pairs was detected as parent-offspring or siblings with phi (Fig S9C and S9D). Among them, four pairs of an NSO and a hybrid in the cluster show almost no zero IBS sites (Fig S9E), indicating they are parent-offspring pairs, while all pairs of a CSO and a hybrid showed a significantly larger proportion of zero IBS sites than 0 (Fig S9F). As we explained above, phi values between SO and hybrids are possibly inflated, so we examined the clusters with high phi value (>0.2) in details with the number of segregating sites (S) and the number of sites where no alleles are shared between two individuals in a pair ($N_{AAaa}$). We found that among the 88 pairs between NSO and hybrid, all the 64 pairs involving an F1 hybrid clustered at IBS0 <0.1 (Fig S9C, S9E, Fig S10), reflecting their low genetic diversity. All the pairs except for the four parent-offspring pairs showed only the average level of S and $N_{AAaa}$ (Fig S10), suggesting these 60 pairs are not related. It is also supported by the fact that backcrosses are not involved in these 60 pairs because non-parent-offspring related pairs should involve backcrosses, not F1 hybrids. We concluded that the high kinship coefficient values of these 60 pairs are inflated by the violation of Hardy-Weinberg equilibrium. Similary, among the 33 pairs between CSO and hybrid, all the 24 pairs involving an F1 hybrid clustered at IBS0 <0.1 (Fig S9D, Fig S9F, Fig S10). We examined these pairs and showed none of them are related (Fig S10). In total, we identified 8 parent-offspring pairs involving 4 different parents and 8 offspring, one pair of full siblings, and one pair of closely related individuals, possibly half siblings (Table S7).

#### 3. mtDNA analyses

The work of Barrowclough et al. 2011 has shown that there are two distinct mitochondrial haplotypes with different geographical distribution in barred owls, using 121 mitochondrial control-region sequences (518 bp) sampled form 18 populations distributed across the United States and Canada. They suggested two populations in the past, possibly two Pleistocene refugia, one was located on the Atlantic Coast and the other was in the south-central part of United States. The two ancient populations represented by these mitochondrial haplotypes have been spread and merged around the boundary areas probably after the last ice age. Barrowclough et al. 2011 also reported that both of the haplotypes were found on the west coast of the United States and in the boundary area, ranging from Minnesota, Michigan, Ohio to Florida. Although we don’t know how long the two populations were separated from each other, nor when they started to merge, the autosomal genetic diversity harbored by the two populations should have been rigorously mixed by recombination since the two populations encountered each other in the boundary area and spreading to the entire barred owl population with gradient proportions.

To confirm that our data contain the genetic components from both of the two ancient populations, we examined haplotypes of mtDNA in our data.

3-1. Extraction of mitochondrial variants

To obtain variants on mitochondrial DNA (mtDNA), we filtered the masked file separately from the nuclear variants using the GATK SelectVariants tool with the "--restrictAllelesTo BIALLELIC --excludeFiltered" options, and then extracted the variants mapped to the mitochondirial genome sequence in the reference file using vcftools(Danecek et al. 2011). We filtered the resulting file and eliminated the sites with the minimum quality of assigned genotype (GQ) smaller than 40. At each site, we took an allele with higher read depth between reference and alternative, and used the alleles with higher read depth than 20.

3-2. Phylogenetic analysis of the mitochondrial non-coding region

In previous works, the mitochondrial control region was used for studies of the phylogeography of spotted owls and barred owls (Haig, Mullins, Forsman, et al. 2004; Barrowclough et al. 2005; Barrowclough et al. 2011), but Hanna et al. (2017)(Hanna et al. 2017) has elucidated that mitochondria of spotted owls and barred owls have duplicated control regions. Because NGS short reads techniques don't give the best quality data for duplicated or repetitive sequences, and variants on them are quite often mapped to wrong copies, we decided to remove the control regions and the genes between them (Figure S11). Instead, we used all the remaining non-control region of mtDNA (15kb) for the phylogenetic analysis with MEGA(Kumar et al. 2016). After removing all the sites with missing individual data, the remaining 327 variants on mtDNA for 51 male and female samples were used. Evolutionary histories were inferred using the Neighbor-Joining method (Saitou and Nei 1987), and the evolutionary distances were computed using the number of differences method (Nei and Kumar 2000) and are in the units of the number of base differences per sequence.

The resulting phylogenetic tree of the mitochondrial non-control region in our data showed clear clusters of SO and BO (Figure S12). In the barred owl cluster, two clades were formed and both of them involve EBO and WBO, supported by higher bootstrap values of 99 and 95. The nucleotide diversity and Hudson's Fst between these two haplotypes on mtDNA were 0.0047 and 0.547 respectively. In the SO clade, two of the three CSO samples, UWBM62061 and ZRHG104, showed the deepest split from the rest, but the other CSO individual, ZRHG103, clustered together with NSO samples (Figure S12). Considering that ZRHG103 is from a hybrid zone between CSO and NSO, in Nevada County in California, it suggests that the individual carries an introgressed mtDNA from NSO, though we need more samples to examine it. Previous works based on mtDNA have reported the presence of CSO haplotypes in the range of NSO, and vice versa (Fleischer et al. 2004; Haig et al. 2004; Barrowclough et al. 2005), and the work on microsatellite loci showed that both long-distance dispersal and hybridization are occurring (Funk et al. 2008). Our result of this mtDNA analysis together with the ancestry analysis of hybrids supports hybridization between CSO and NSO.

3-3. Identification of mitochondrial lineages

To identify the haplotypes reported by Barrowclough et al. 2011 in our data, we identified five fixed differences between “Atlantic Coast” and “south-central” haplotypes using their 121 mitochondrial control-region sequences (518 bp) (JN097839 – JN098025) with MEGA software. Among the five SNPs, we found two (position 14886 and 14947) in our vcf file, while the other three were filtered out. We checked the genotypes of 51 samples at these two sites in our data, and found that at position 14886, T and A are segregating in our data, while G and A are segregating in the 121 sequences, suggesting mis-mapping of reads. At position 14947, T and C are segregating in both sets of data. We identified the two mitochondrial haplotypes in Barrowclough et al. 2011 on our non-control region sequences with position 14947 (Figure S12). We identified 24 and 14 sequences as linked sequences to the Atlantic Coast and the south-central haplotypes of the control region, respectively. Although this identification depends on a single variant, these two haplotypes corresponded perfectly to the two clusters on the phylogenetic tree (Figure S12). The geographic distribution of these haplotypes was quite similar to the one shown using the control region (Fig S13 and Figure 1 in Barrowclough et al. 2011). These results revealed that our samples include the known population structure observed on mtDNA, suggesting that our data also contains the autosomal genetic variety accumulated in both of the two ancient populations at least partially. The genetic varieties accumulated in the two distinct ancient populations found in an individual might be the cause of the complicated SMC++ patterns for barred owls.

**Captions for Supplementary Figures**

Figure S1. Comparison of the distribution of the lengths of scaffolds and contigs between the assemblies.

The improved contiguity of our new assembly (StOcCau_2.0) was shown in comparison with the previous one (StOcCau_1.0).

Figure S2. Histogram of the mean read depth of scaffolds (>1Mb) in males and females.
We calculated the averaged read depth for each scaffold across variants and individuals for males and females.

Figure S3. Histogram of the proportion of missing data in scaffolds and contigs (<1Mb, >=100kb) in males and females.
The mean proportion of missing data was calculated across individuals for each scaffold or contig (<1Mb, >=100kb), separately for males and females.

Figure S4. Description of variants identified on autosomes and sex chromosomes.
A-C. The mean proportion of missing data for each scaffold/contig was plotted for males against that in females for autosomes (A), the Z (B) and the W chromosome (C).
D-F. The number of genotypes was plotted against the length of the scaffold/contig for males (blue) and females (orange), and for autosomes (D), the Z (E) and the W chromosomes (F).

Figure S5. PCA plot for barred owls.
PCA plot for WBO and EBO samples only was shown with sample names.

Figure S6. Comparison of pi and $F_{ST}$ among chromosome types.
Nucleotide diversity (pi) within (A) and between (B) populations and Hudson's $F_{ST}$ (C) were shown for autosomes and the Z and W chromosomes.

Figure S7. Comparison of two estimators of the kinship coefficient.
The two estimations of the kinship coefficient (phi) calculated with Equation(9) and Equation(11) from Manichaikul et al. 2010(Manichaikul et al. 2010) were compared. The two estimations would be identical (on the diagonal line) if there is no violation of HWE. Two values were compared for pairs of samples within

EBO (A), WBO (B), NSO (C), CSO (D) and hybrids (E).

Figure S8. Inference of related individuals within populations.
Phi values are plotted against the proportion of zero IBS (the portion of the sites where two individuals share no alleles identical by state) for each pair of samples within NSO (A), CSO (B), EBO (C) and WBO (D). Dashed lines are inference criteria of phi (from Table 1 in Manichaikul et al. 2010(Manichaikul et al. 2010)) for parent-offspring pairs and full siblings (PO + FS) and $2^{nd}$ degree relations (2D).

Figure S9. Inference of related individuals between populations.
Phi values are plotted against the proportion of zero IBS (the portion of the sites where two individuals share no alleles identical by state) for each pair of samples involving EBO and hybrids (A), WBO and hybrids (B), NSO and hybrids (C), and CSO and hybrids (D). The squared parts in (C) and (D) were enlarged in (E) and (F) respectively. Dashed lines are inference criteria of phi (from Table 1 in Manichaikul et al. 2010(Manichaikul et al. 2010)) for parent-offspring pairs and full siblings (PO + FS) and $2^{nd}$ degree relations (2D).

FigureS10. Sampling location, number of segregating sites and number of zero IBS sites for the pairs with high phi values.
Sampling location, number of segregating sites (S) and number of zero IBS sites ($N_{AAaa}$) were shown for the pairs with high phi and low IBS0 values involving a hybrid and a SO sample. All the 64 pairs of samples in FigS7 (E) and all the 24 pairs in the FigS7 (F) are shown in green. Among them, four parent-offspring pairs detected with IBS0 values are shown in pink. All the individuals involved in the parent-offspring pairs are from Humboldt County in California (shown in orange). The remaining sympatric pairs showed only the average level of S and $N_{AAaa}$.

Figure S11. The mean DP and the number of missing individual data for variants on mtDNA.

The mean read depth (blue) and the number of missing individual data (orange) were shown. The region containing the duplicated control regions, two tRNA genes and the ND6 gene (position 14879~) was removed from the phylogenetic analysis.

Figure S12. The phylogenetic tree on the non-control region of mtDNA

Variants on the non-control region (15kb) of all the 51 samples were involved. The sum of the branch length was 337.19. After eliminating all positions containing gaps and missing data, there were a total of 327 positions in the final dataset. The evolutionary history was inferred using the Neighbor-Joining method. The percentages of replicate trees in which the associated taxa clustered together in the bootstrap test (1000 replicates) are shown when it's higher than 50%. Atlantic Coast and the south-central haplotypes were identified using a SNP in a control region.

Figure S 13. Geographic distribution of the mitochondrial haplotypes.
Distribution of the Atlantic Coast (blue, 24 individuals) and the south-central (magenta, 14 individuals) haplotypes in our data were shown to be compared with Figure 1 in Barrowclough et al. 2011(Barrowclough et al. 2011). The size of circles and pie charts correspond to the number of samples.

**Captions for Supplementary Tables**

TableS1. Metrics of Assemblies.
Summary statistics for our new assembly (StrOccCau_2.0) in comparison with the previous assembly (StrOccCau_1.0). We removed contigs and scaffolds shorter than 1 kb from our assembly before calculating these statistics to make it comparable to StrOccCau_1.0. N50, L50, the percentage of missing data (Ns) and the total length of scaffolds and contigs are shown together with the number of scaffolds and contigs longer than 1kb or 1Mb.

TableS2. Sample information.
Short sample IDs used in this study were shown with corresponding museum specimen IDs. All samples were primarily identified by morphology ("Primary

identification”), then re-identified using the genetic data (“Genetic identification”). Genetically identified sex (M; Male, F; Female), sampling locations, and the mean and the standard deviation of read depth across variants were shown. The mean and the standard deviation of read depth were calculated for variants on scaffolds longer than 1Mb after the basic filtering. The averaged read depth across the sample means was 31.7 and the standard deviation was 6.51. The column “SRA ACCN” provides NCBI Sequence Read Archive (SRA) run accessions in which the raw sequences for each sample are archived.

TableS3. Detail description of the 82 scaffolds identified on autosomes.
The mean read depth (DP), the mean number of variants, and the mean portion of missing data across samples for each of 82 scaffolds identified on autosomes were shown with their standard deviations. The variants from the filtered set of SNPs, after removing individual variants with GQ smaller than 40, were used for this table (as used in the diversity analysis). The values for the total 51 samples, and for males and females were shown and no significant difference between sexes were there.

TableS4. Detail description of the 15 scaffolds identified on the Z chromosome.
The mean portion of missing data, the mean read depth (DP), and the mean number of variants across samples for each of the 15 scaffolds identified on the Z chromosome were shown with their standard deviations for males and females. The variants from the filtered set of SNPs, after removing individual variants with GQ smaller than 40, were used to make the table comparable with Table S3.

TableS5. Detail description of the 44 scaffolds/contigs identified on the W chromosome.
The mean portion of missing data, the mean read depth (DP), and the mean number of variants across samples for each of the 44 scaffolds and contigs identified on the W chromosome were shown with their standard deviations for males and females. The variants from the filtered set of SNPs, after removing individual variants with GQ smaller than 40, were used to make the table comparable with Table S3.

TableS6. Ancestral components of hybrid samples.

A. The portion of “spotted owl alleles” in hybrids at the apparent fixed differences between SO and BO was shown. Heterozygosity at these sites and inferred status of hybrids are shown.

B. The portion of “NSO allele” and “CSO allele” at the apparent fixed differences between NSO and CSO, where no polymorphism was observed in BO, was shown. The sum of the two components is 0.5, because these values were calculated under the assumption that one of the parents of a hybrid is a barred owl.

TableS7. List of closely related samples.

A. Detected pairs of related samples were shown with their phi, the portion of the sites where two individuals share no alleles identical by state (IBS0), and inference of their relationship.

B. Parsimonious list of related samples. The four samples removed from the diversity analyses and demography analyses were marked with asterisks.

TableS8. Nucleotide diversity within and between populations on autosomal variants. A set of variants after filtering out the individual variants with GQ smaller than 40 was used.

Table S9. Weir and Cockerham’s $F_{ST}$ for each pair of populations.
Weir and Cockerham‘s weighted $F_{ST}$ for each pair of populations was calculated using autosomal variants.

Table S10. The number of segregating sites and Tajima’s D.
The number of samples and the number of segregating variants in two different sets of variants were shown for each population. The set retaining all the individual variants with GQ greater than 40 was used for the diversity analysis, and the set retaining only the variants with no missing data was used for all the other analyses. The four WBO individuals those have closely related individuals in the sample set were removed here. Tajima’s D, and the averaged number of

variants across the windows used to calculate Tajima's D were also shown.

Table S11. Female ancestry of hybrids.
The proportion of "spotted owl alleles" and the inferred ancestry of the W chromosome were shown.

Table S12. The number of private alleles.
The number of private alleles (singletons and private homozygotes) (NP) in groups are shown for BO (A) and SO (B). NP in a test individual in each group (NP_test) and the mean NP for focal individuals (Mean NP_focal) are also shown.

TableS13. Nucleotide diversity within and between populations on the sex chromosomes.

Wilson Sayres MA, Lohmueller KE, Nielsen R. 2014. Natural Selection Reduced Diversity on Human Y Chromosomes. PLoS Genet. 10.

### Supplementary figures

Fig. S1 Comparison of the distribution of the lengths of scaffolds and contigs between the assemblies.

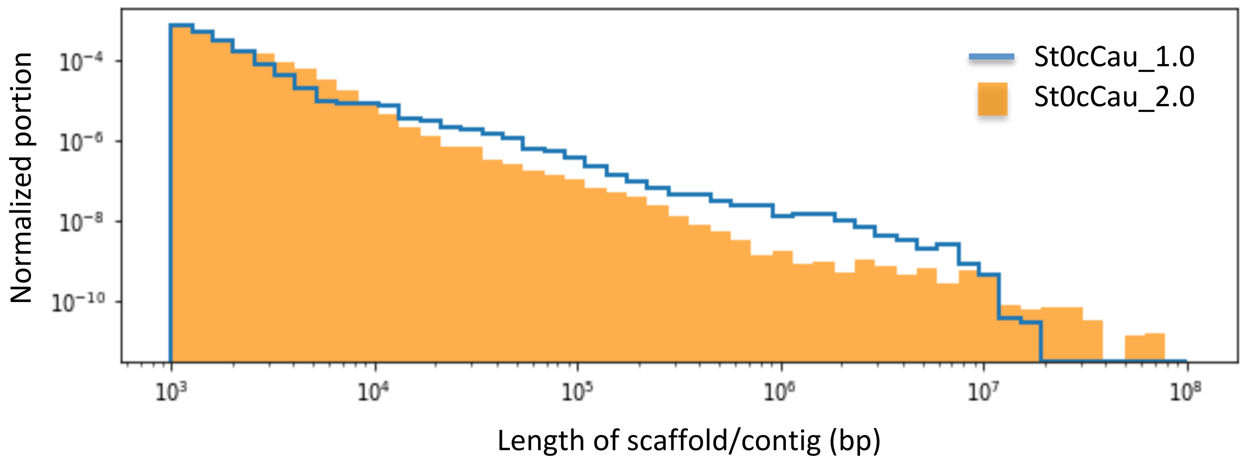

Fig. S2 Histogram of the mean read depth of scaffolds (>= 1 Mb) in males and females.

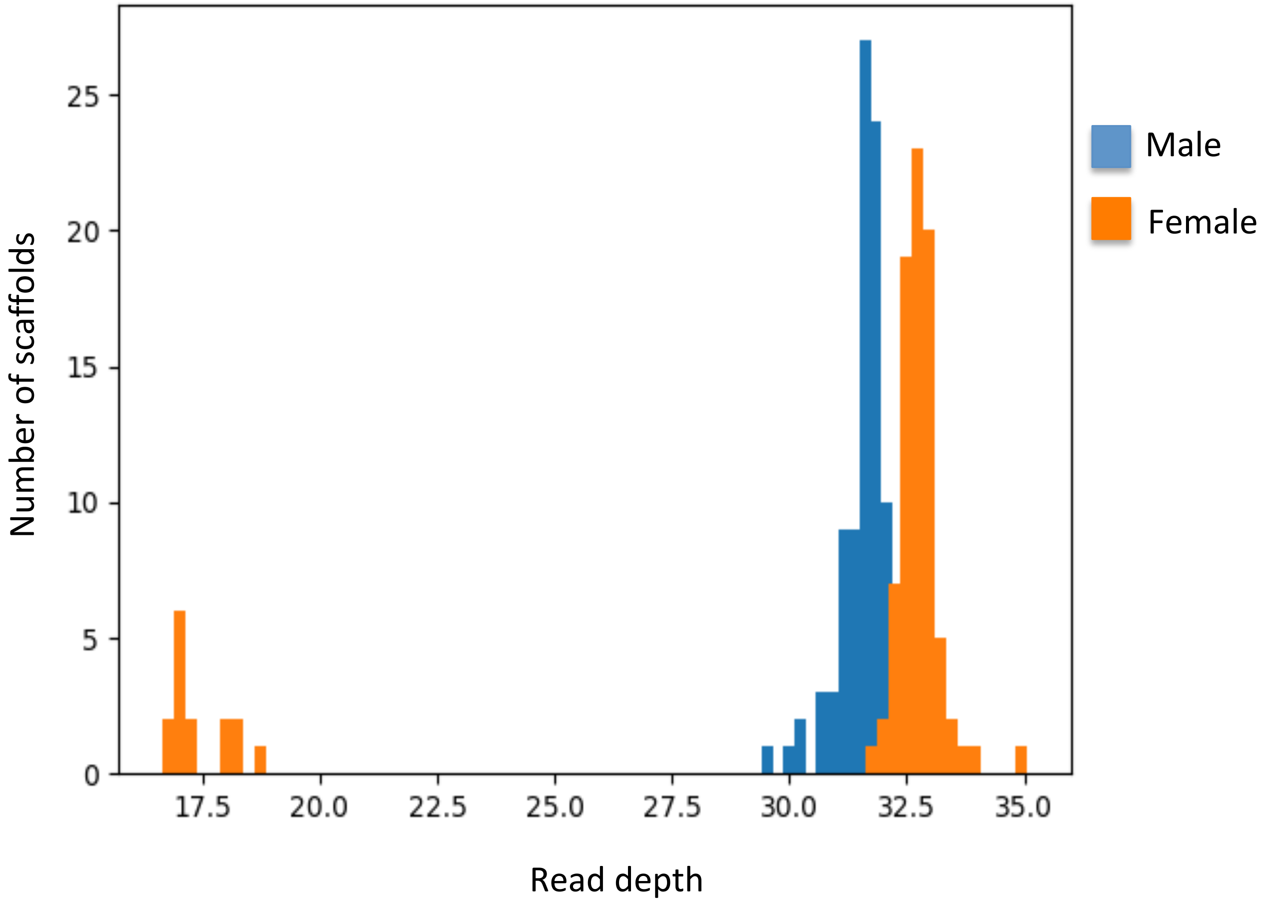

Fig. S3 Histogram of the proportion of missing data in scaffolds and contigs (>= 100kb, < 1Mb) in males and females.

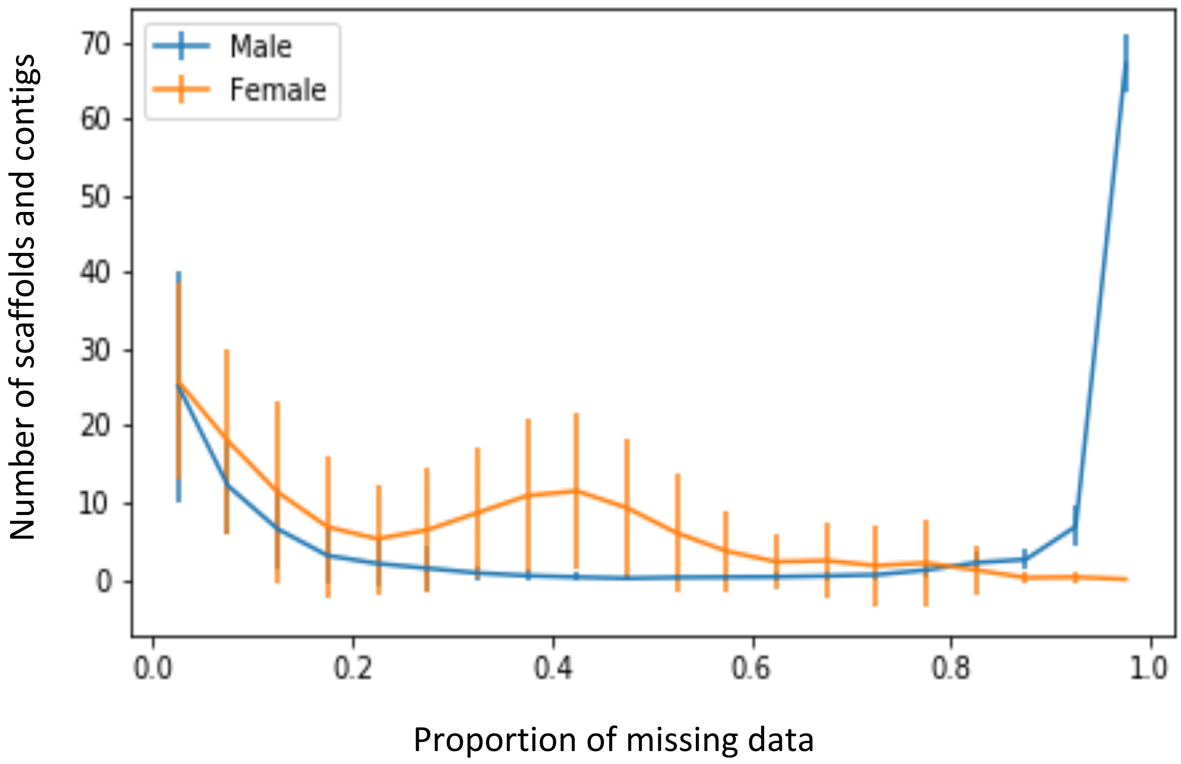

Fig. S4

Description of variants identified on autosomes and sex chromosomes.

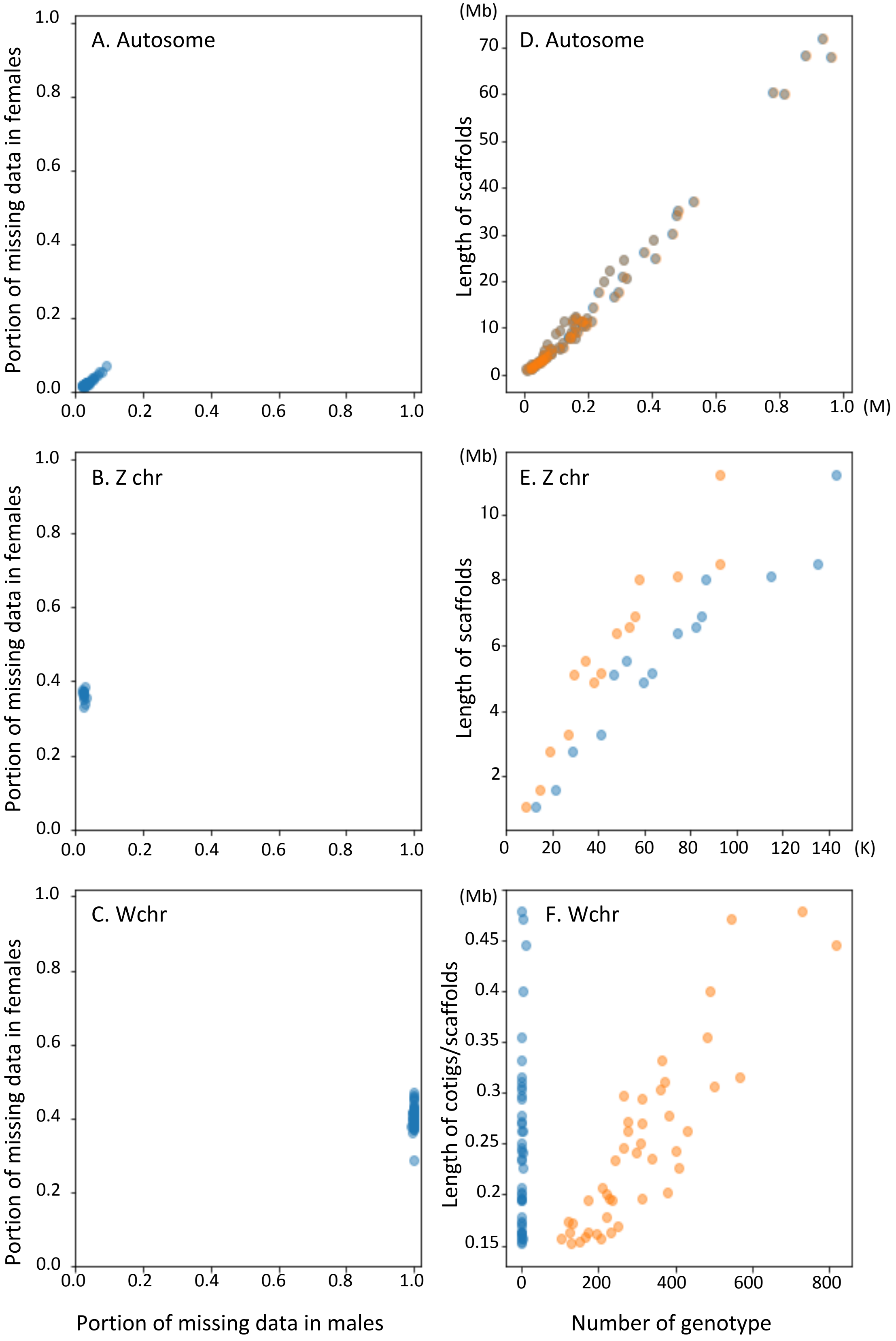

Fig. S5 PCA plot of Barred Owls.

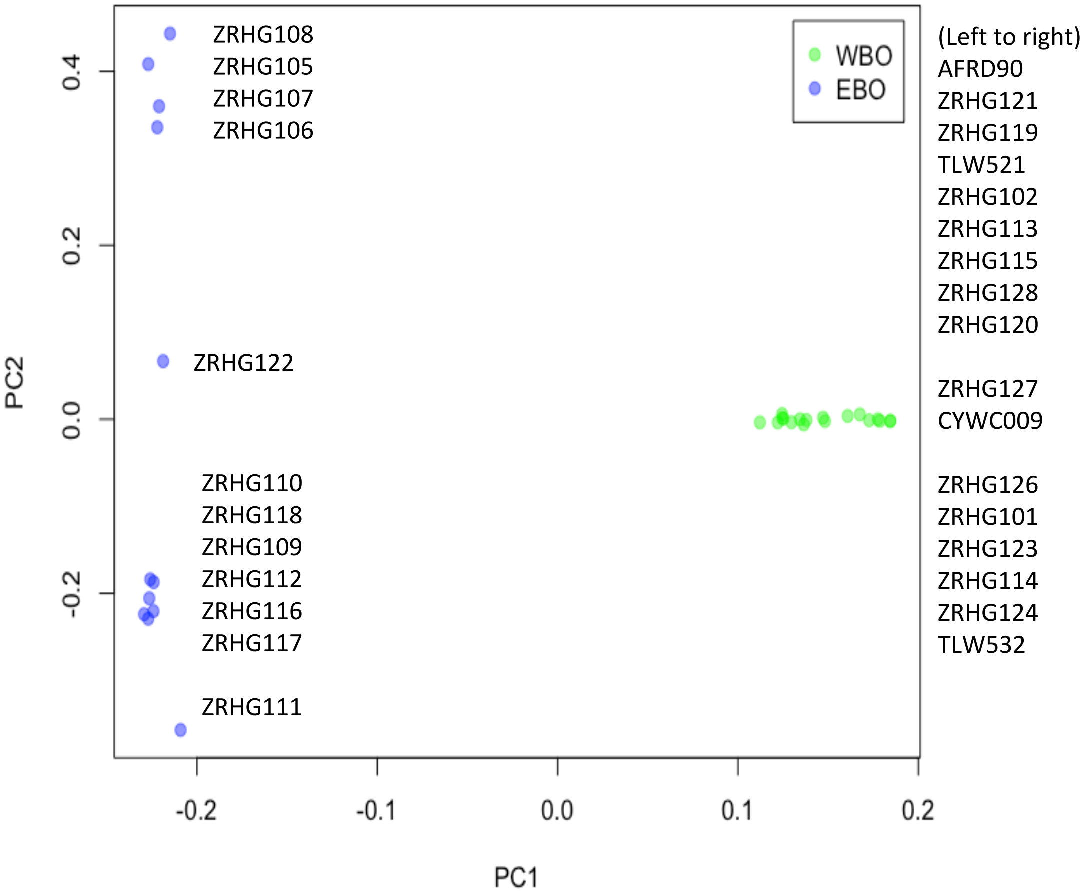

Fig. S6 Comparison of pi and $F_{ST}$ among chromosome types.

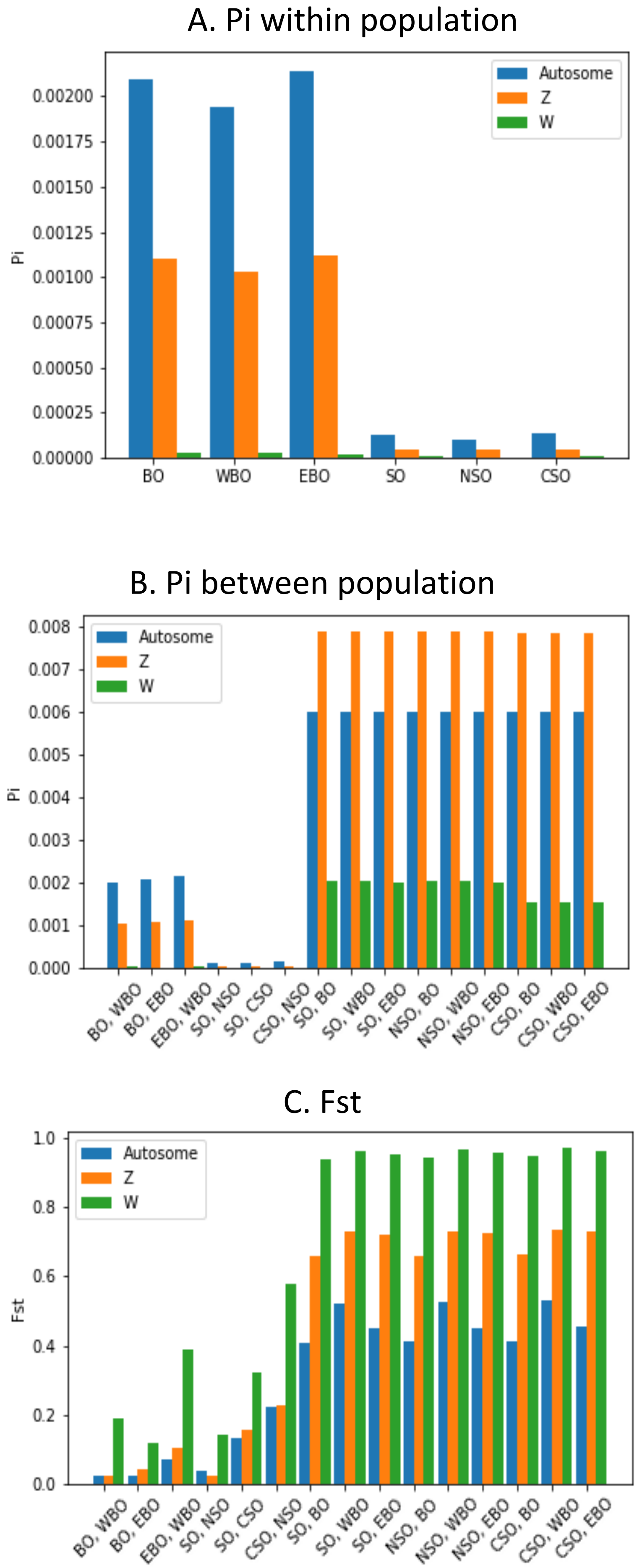

Fig. S7 Comparison of the two estimators of the kinship coefficient.

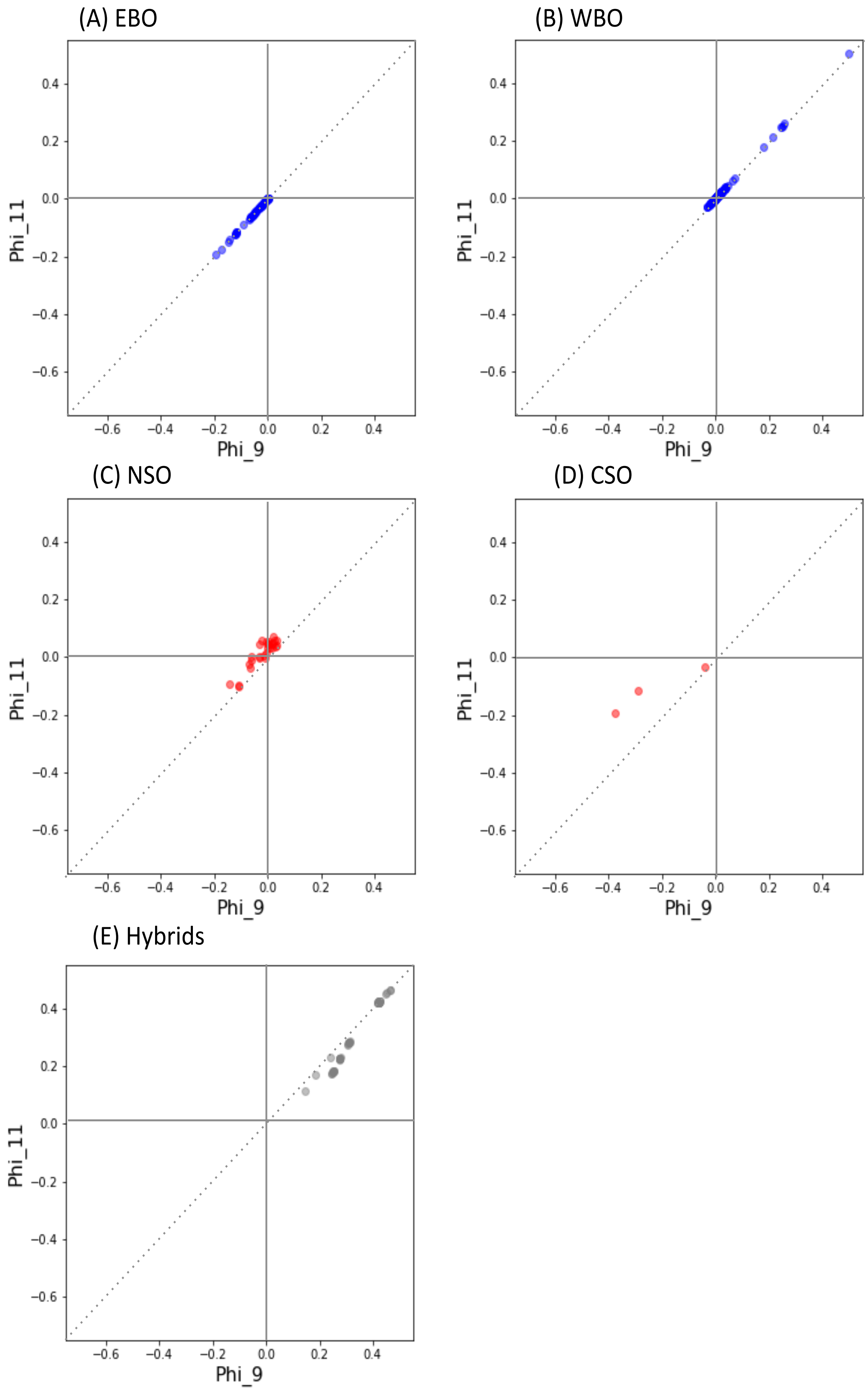

Fig. S8 Inference of related individuals within populations.

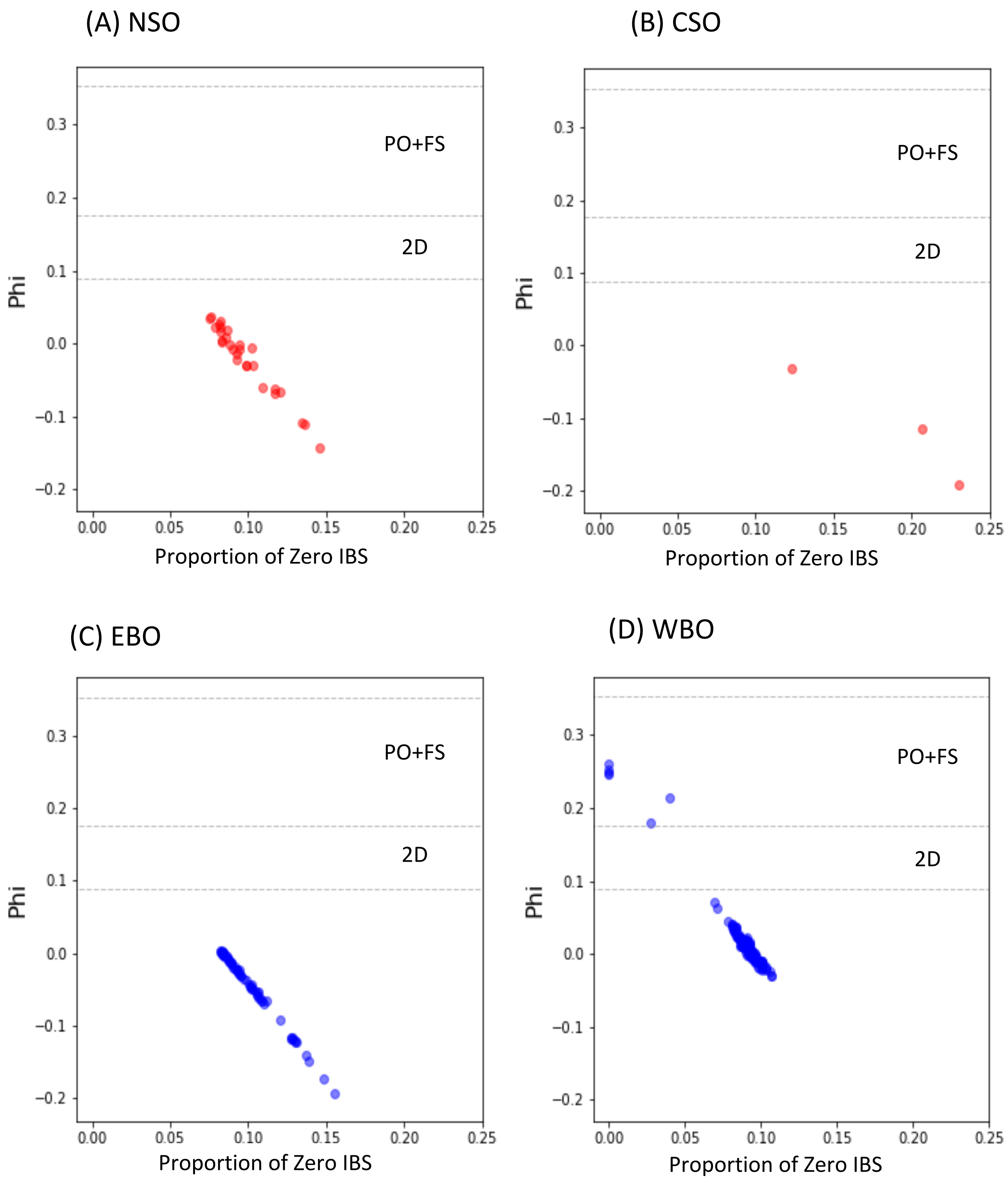

Fig. S9 Inference of related individuals between populations.

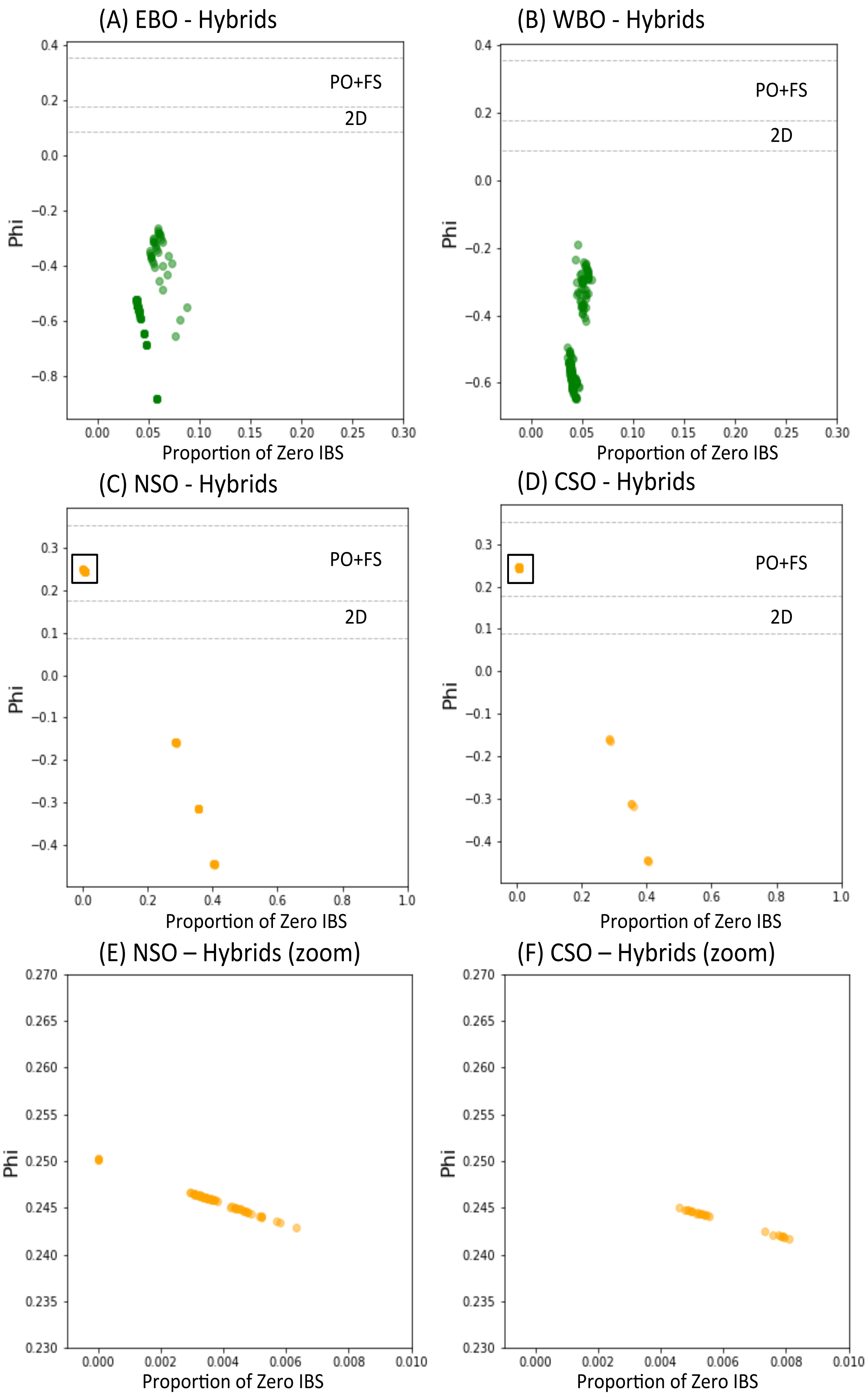

### Fig.S10 Sampling location, number of segregating sites and number of zero IBS sites for the pairs with high phi values.

| ID | Category1 | Category2 | Sampling location: County | Sampling location: State |
|---|---|---|---|---|
| Sequoia | SO | NSO | Marin County | California |
| 2018-365 | SO | NSO | Marin County | California |
| ZRHG125 | SO | NSO | Humboldt County | California |
| ZRH615 | SO | NSO | Humboldt County | California |
| ZRH625 | SO | NSO | Humboldt County | California |
| UWBM48266 | SO | NSO | Whatcom County | Washington |
| UWBM91379 | SO | NSO | Linn County | Oregon |
| UWBM91381 | SO | NSO | Baker County | Oregon |
| ZRHG103 | SO | CSO | Nevada County | California |
| ZRHG104 | SO | CSO | San Diego County | California |
| UWBM62061 | SO | CSO | Tulare County | California |
| 1957-00137 | Hybrid | F1 | Humboldt County | California |
| 1957-00240 | Hybrid | F1 | Humboldt County | California |
| 1957-00243 | Hybrid | F1 | Humboldt County | California |
| ZRH600 | Hybrid | F1 | Humboldt County | California |
| ZRH610 | Hybrid | F1 | Humboldt County | California |
| ZRH962 | Hybrid | F1 | Humboldt County | California |
| LCW1363 | Hybrid | F1 | Shasta County | California |
| LCW1383 | Hybrid | F1 | Siskiyou County | California |
| TLW519 | Hybrid | Backcross | Lane County | Oregon |
| TLW528 | Hybrid | Backcross | Benton County | Oregon |
| ZRH607 | Hybrid | Backcross | Benton County | Oregon |

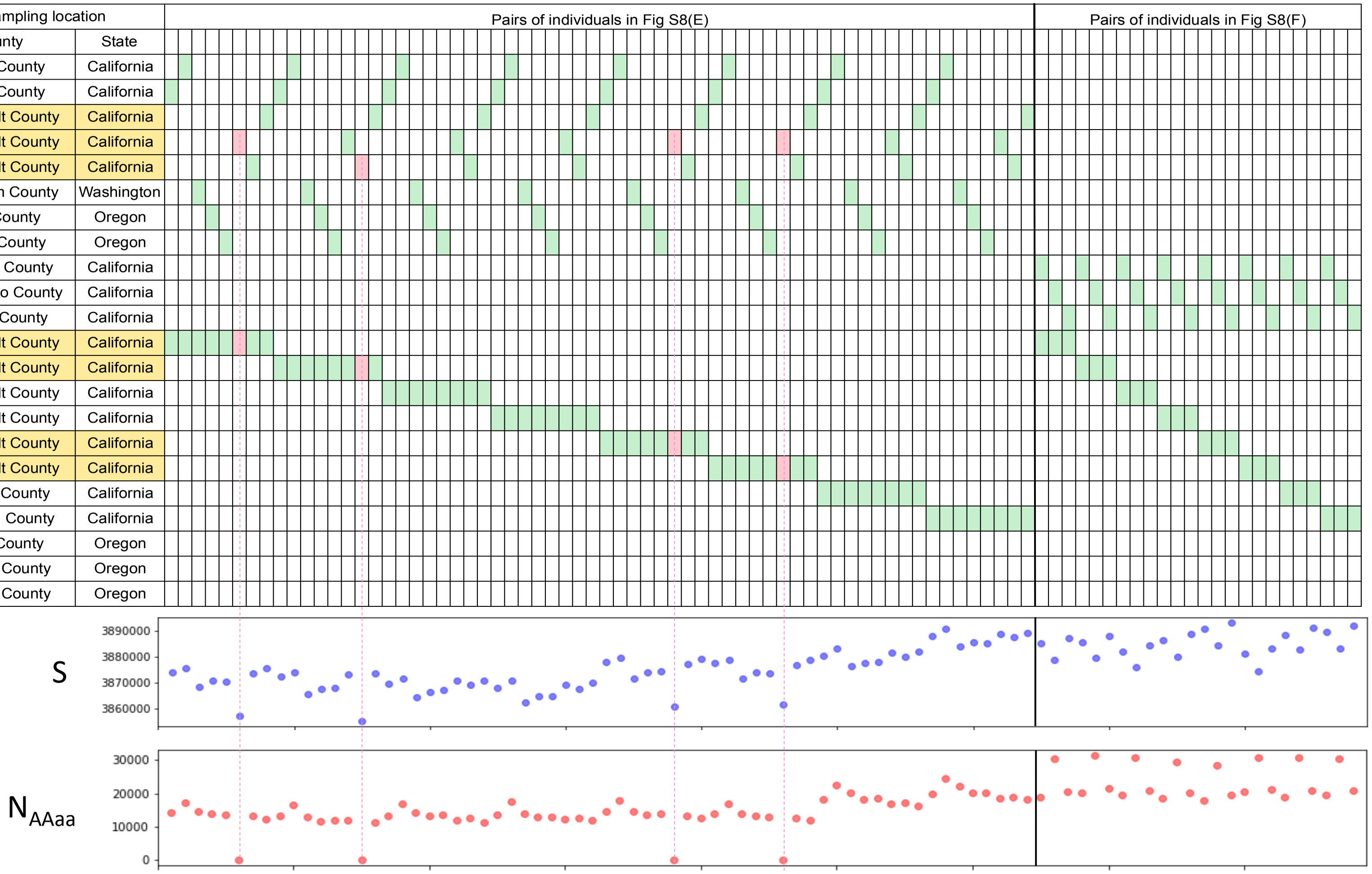

Fig. S11 The mean DP and the number of missing individual data for variants on mtDNA.

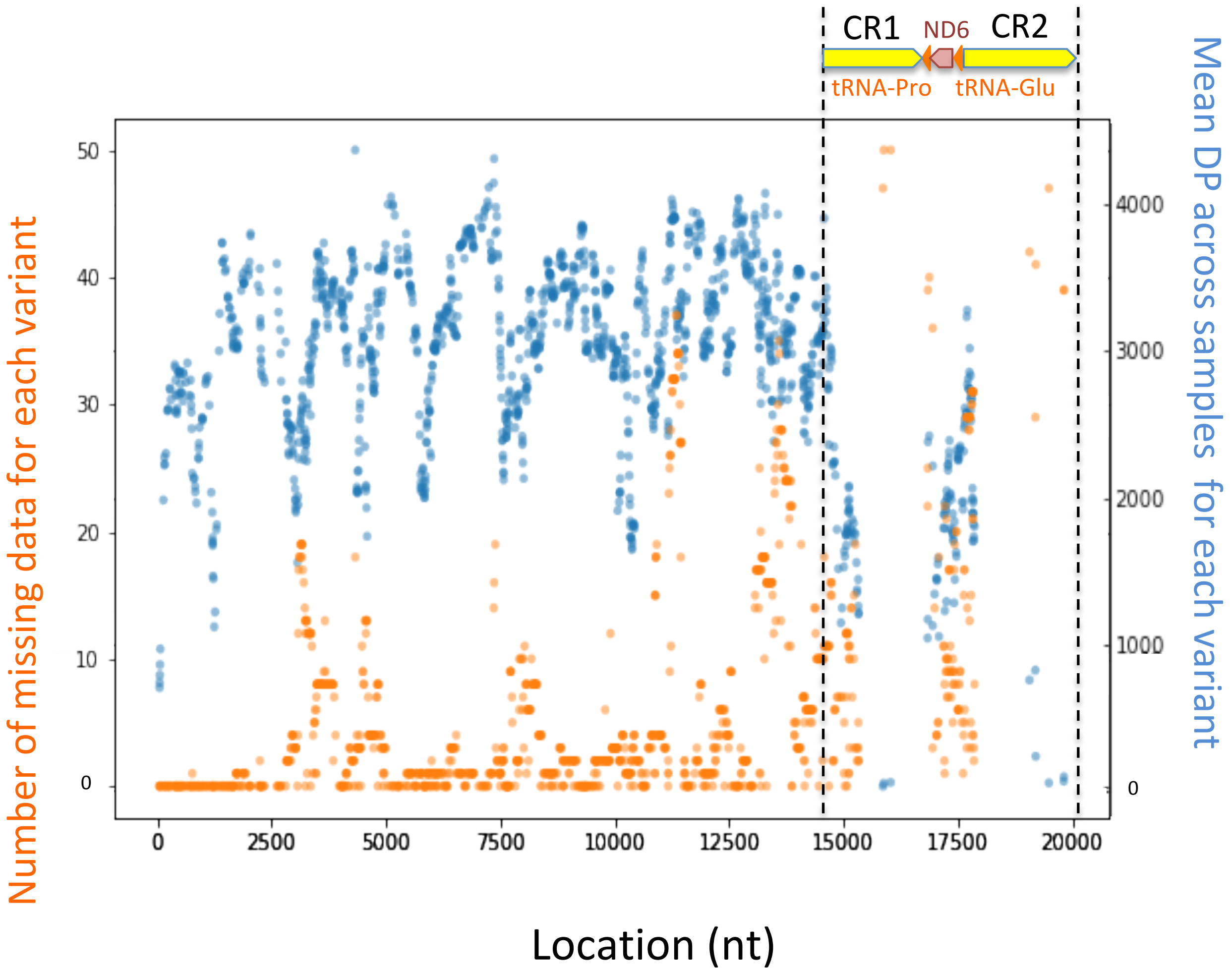

Fig. S12 The phylogenetic tree on the non-control region of mtDNA.

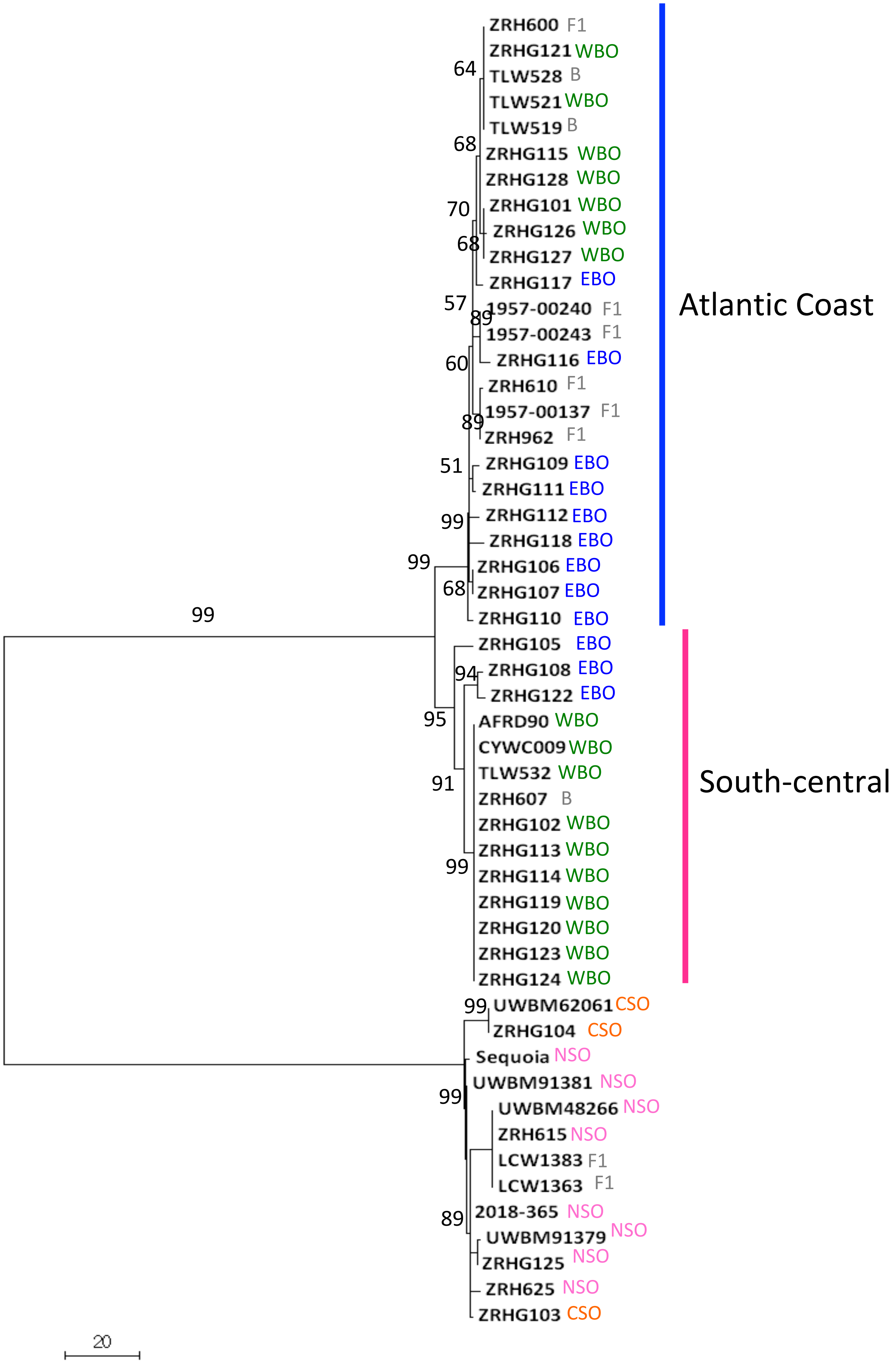

Fig. S13 Geographic distribution of the mitochondrial haplotypes.

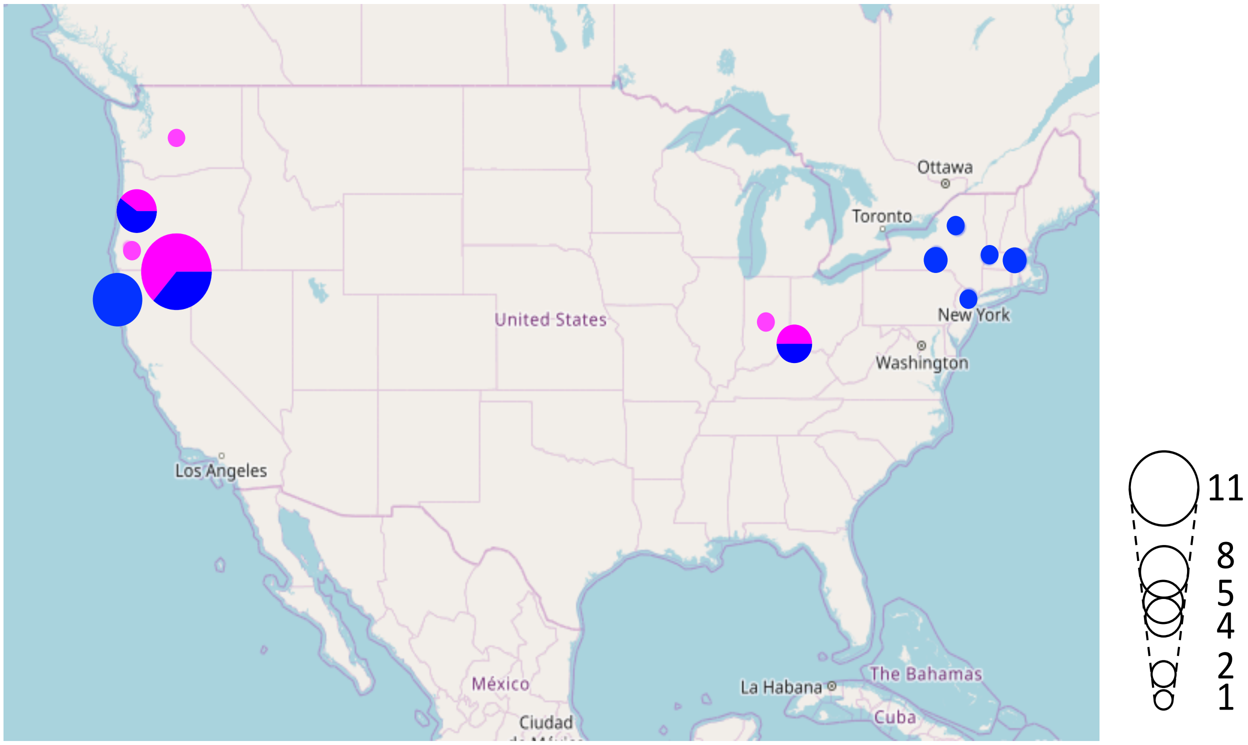

**Supplementary tables**

TableS1. Metrics of Assemblies.

| Assembly | N50 (kb) | L50 | Ns (%) | Total length of scf/ctg >1kb (Mb) | Number of scf/ctg > 1kb | Total length of scf/ctg >1Mb (Mb) | Number of scf/ctg > 1Mb |
|---|---|---|---|---|---|---|---|
| StrOccCau_1.0 | 3,983.0 | 92 | 1.10 | 1,255.5 | 8108 | 1,075.5 | 303 |
| StrOccCau_2.0 | 20,549.8 | 16 | 1.91 | 1,254.4 | 11568 | 1,173.7 | 97 |

Table S6. Ancestral components of hybrid samples.

A. Portion of SO ancestry in hybrids

| ID | Portion of SO ancestry | Heterozygosity | State |
|---|---|---|---|
| 1957-00137 | 0.500 | 1.000 | F1 |
| 1957-00240 | 0.500 | 1.000 | F1 |
| ZRH610 | 0.500 | 1.000 | F1 |
| ZRH962 | 0.500 | 1.000 | F1 |
| ZRH600 | 0.500 | 1.000 | F1 |
| 1957-00243 | 0.500 | 1.000 | F1 |
| LCW1363 | 0.500 | 0.999 | F1 |
| LCW1383 | 0.500 | 0.999 | F1 |
| TLW519 | 0.323 | 0.646 | F1 x BO |
| ZRH607 | 0.276 | 0.552 | F1 x BO |
| TLW528 | 0.240 | 0.481 | F1 x BO |

B. Portion of NOS/CSO ancestries in hybrids

| State | ID | Portion of NSO ancestry | Portion of CSO ancestry | Total |
|---|---|---|---|---|
| **F1** | 1957-00137 | 0.500 | 0.000 | 0.500 |
| | 1957-00240 | 0.500 | 0.000 | 0.500 |
| | ZRH610 | 0.500 | 0.000 | 0.500 |
| | ZRH962 | 0.500 | 0.000 | 0.500 |
| | ZRH600 | 0.480 | 0.020 | 0.500 |
| | 1957-00243 | 0.466 | 0.034 | 0.500 |
| | LCW1363 | 0.256 | 0.244 | 0.500 |
| | LCW1383 | 0.227 | 0.273 | 0.500 |
| **F1 x BO** | TLW519 | 0.380 | 0.120 | 0.500 |
| | ZRH607 | 0.380 | 0.120 | 0.500 |
| | TLW528 | 0.282 | 0.218 | 0.500 |

Table S7. List of closely related samples.

A. Full list of closely related pairs.

| | Individual 1 | Individual 2 | Phi | IBS0 | Inference |
|---|---|---|---|---|---|
| Within BO | ZRHG101 | ZRHG119 | 0.248 | 5.73E-06 | Parent - offspring |
| | ZRHG101 | ZRHG126 | 0.246 | 8.10E-06 | Parent - offspring |
| | ZRHG114 | ZRHG123 | 0.213 | 4.02E-02 | Full siblings |
| | ZRHG114 | ZRHG124 | 0.252 | 1.75E-05 | Parent - offspring |
| | ZRHG123 | ZRHG124 | 0.260 | 1.37E-05 | Parent - offspring |
| | ZRHG126 | ZRHG127 | 0.179 | 2.76E-02 | Half siblings |
| Hybrid - SO | 1957-00137 | ZRH615 | 0.250 | 6.48E-06 | Parent - offspring |
| | 1957-00240 | ZRH625 | 0.250 | 5.45E-06 | Parent - offspring |
| | ZRH610 | ZRH615 | 0.250 | 1.22E-05 | Parent - offspring |
| | ZRH962 | ZRH615 | 0.250 | 7.25E-06 | Parent - offspring |

B. Parsimonious list of related samples.

| ID | Population | Related individuals |
|---|---|---|
| ZRH615 | NSO | Father of ZRH610 (F1), 1957-00137 (F1) and ZRH962 (F1) |
| ZRH625 | NSO | Father of 1957-00240 (F1) |
| ZRHG101* | WBO | Offspring of ZRHG119 (WBO) and ZRHG126 (WBO) |
| ZRHG124* | WBO | Parent or offspring of ZRHG114 (WBO) and ZRHG123 (WBO) |
| ZRHG123* | WBO | Full sibling of ZRHG114 (WBO) |
| ZRHG127* | WBO | Closely related samples (potentially half-siblings) of ZRHG126 (WBO) |

TableS8. Nucleotide diversity within and between populations on autosomal variants.

| Population | SO | NSO | CSO | BO | WBO | EBO |
|---|---|---|---|---|---|---|
| SO | 1.27E-04 | | | | | |
| NSO | 1.12E-04 | 1.03E-04 | | | | |
| CSO | 1.42E-04 | 1.53E-04 | 1.34E-04 | | | |
| BO | 6.02E-03 | 6.02E-03 | 6.01E-03 | 2.10E-03 | | |
| WBO | 6.01E-03 | 6.01E-03 | 6.00E-03 | 2.00E-03 | 1.94E-03 | |
| EBO | 6.01E-03 | 6.01E-03 | 6.00E-03 | 2.10E-03 | 2.14E-03 | 2.14E-03 |

TableS9. Weir and Cockerham's $F_{ST}$ for each pair of populations.

| Pop1 | Pop2 | Weighted $F_{ST}$ |
|---|---|---|
| SO | BO | 0.765 |
| SO | WBO | 0.818 |
| SO | EBO | 0.806 |
| NSO | CSO | 0.253 |
| NSO | BO | 0.750 |
| NSO | WBO | 0.799 |
| NSO | EBO | 0.785 |
| CSO | BO | 0.713 |
| CSO | WBO | 0.748 |
| CSO | EBO | 0.727 |
| EBO | WBO | 0.050 |

TableS10. The number of segregating sites and Tajima’s D.

| Population | Number of samples | Number of segregating variatns (GQ>=40) | Number of segregating variants (GQ>=40) without missing data | Tajima'sD (Std) | Mean number of variants in a100kb window (Std) |
|---|---|---|---|---|---|
| SO | 11 | 539,226 | 456,255 | -0.470 (1.055) | 41.8 (24.6) |
| CSO | 3 | 312,929 | 300,698 | 0.146 (1.094) | 27.5 (24.6) |
| NSO | 8 | 409,928 | 360,001 | -0.634 (1.028) | 32.9 (23.6) |
| BO | 25 | 11,360,737 | 8,572,962 | -0.351 (0.284) | 784.6 (278.6) |
| WBO | 13 | 7,654,945 | 6,474,001 | 0.211 (0.349) | 592.5 (224.7) |
| EBO | 12 | 9,810,520 | 8,550,852 | -0.516 (0.254) | 782.5 (285.1) |

TableS11. Female ancestry of hybrids.

| State | ID | SO ancestry on W chr | Origin of W chr |
|---|---|---|---|
| F1 | LCW1363 | 100.00% | SO |
| | LCW1383 | 100.00% | SO |
| | 1957-00240 | 0.00% | BO |
| | ZRH962 | 0.12% | BO |
| F1 x BO | TLW519 | 0.00% | BO |
| | ZRH607 | 0.00% | BO |

Table S13. Nucleotide diversity within and between populations on the sex chromosomes.

A. Z chromosome

| | SO | NSO | CSO | BO | WBO | EBO |
|---|---|---|---|---|---|---|
| SO | 5.02E-05 | | | | | |
| NSO | 4.39E-05 | 4.40E-05 | | | | |
| CSO | 5.46E-05 | 5.99E-05 | 4.92E-05 | | | |
| BO | 7.89E-03 | 7.89E-03 | 7.87E-03 | 1.10E-03 | | |
| WBO | 7.89E-03 | 7.88E-03 | 7.87E-03 | 1.04E-03 | 1.03E-03 | |
| EBO | 7.88E-03 | 7.88E-03 | 7.86E-03 | 1.09E-03 | 1.13E-03 | 1.12E-03 |

B. W chromosome

| | SO | NSO | CSO | BO | WBO | EBO |
|---|---|---|---|---|---|---|
| SO | 9.09E-06 | | | | | |
| NSO | 5.47E-06 | 5.09E-06 | | | | |
| CSO | 6.89E-06 | 8.64E-06 | 6.36E-06 | | | |
| BO | 2.05E-03 | 2.04E-03 | 1.56E-03 | 3.20E-05 | | |
| WBO | 2.03E-03 | 2.03E-03 | 1.55E-03 | 3.04E-05 | 2.85E-05 | |
| EBO | 2.02E-03 | 2.01E-03 | 1.54E-03 | 2.49E-05 | 3.74E-05 | 2.11E-05 |
